# Structural modelling and functional analysis support lipid binding by the dimeric *E. coli* DedA protein YqjA

**DOI:** 10.64898/2026.09.20.752871

**Authors:** Thomas J. Paige, Junxi Chen, Aline Le Roy, Connor D. D. Sampson, Benjamin F. Cooper, Matthew Batson, Hollie L. Scarsbrook, Lucy R. Forrest, Georgia L. Isom, Caroline Mas, Vanessa Leone, Christopher Mulligan

## Abstract

Maintenance of membrane homeostasis is essential for bacterial viability, yet the molecular functions of many membrane proteins involved in this process remain poorly understood. The widely distributed DedA superfamily of integral membrane proteins has been implicated in membrane homeostasis, with deletion of DedA genes resulting in sensitivities to temperature, pH and a range of antimicrobial compounds. Several bacterial DedA proteins have been linked to lipid transport, providing a potential connection between function and phenotype. However, a lack of direct functional and structural data means that the precise role of DedA proteins in bacterial membrane homeostasis remains unclear. Here, using analytical ultracentrifugation (AUC), we show that the *Escherichia coli* DedA protein YqjA exists predominantly as a dimer and, by combining structural modelling with site-specific cysteine crosslinking, we have identified the most likely dimer interface. Modelling of dimeric YqjA in the presence of lipids predicts an interfacial lipid-binding site located in close proximity to several conserved, functionally important residues. Consistent with lipid binding, addition of lipid substantially increased the thermal stability of YqjA. Furthermore, mutation of residues associated with the predicted binding site impaired YqjA function. Together, these findings provide new structural and functional insights into the DedA family and support a role for lipid binding in the activity of YqjA.

## Introduction

Maintaining the integrity of the bacterial cell envelope under a variety of environmental conditions is essential for survival and pathogenesis. Proper membrane homeostasis is required for numerous key membrane-associated processes, including nutrient uptake, signal transduction, cell division and export of toxins and antimicrobial compounds. Maintaining homeostasis requires the coordinated effort of an array of membrane proteins, many of which remain poorly characterised. One family that has recently emerged as an important contributor to membrane homeostasis is the DedA superfamily.

The DedA superfamily is a widely distributed and highly conserved family of integral membrane proteins found throughout the bacterial kingdom, as well as in eukaryotes and archaea^1^. Despite their broad distribution, the physiological and molecular functions of DedA proteins remain poorly understood. However, genetic studies have demonstrated that DedA proteins play important roles in bacterial fitness across a diverse range of species. In *Borrelia burgdorferi*, deletion of the sole DedA gene (*bb0250*) leads to severe cell division defects and eventual cell death^2^, demonstrating members of the DedA family can be essential for cell viability. In addition, studies have linked DedA proteins to a range of membrane-associated phenotypes, including sensitivity to membrane-targeting antimicrobials. DedA homologues have been implicated in resistance to a macrocyclic alkaloid halicyclamine A in *Mycobacterium bovis*^3^. In *Burkholderia thailandensis*, disruption of the DedA gene *dbcA* not only disrupts proton motive force maintenance^4^, but also sensitises the bacterium to the cationic antimicrobial peptide (CAMP) colistin due to the requirement for DbcA in aminoarabinose (Ara4N) modification of Lipid A^4,5^. Consistent with this, DedA proteins have also been implicated in resistance to colistin in other organisms such as *Klebsiella pneumoniae*, *Enterobacter cloacae* and *Burkholderia glumae*^6–8^, as well as CAMP resistance in *Salmonella enterica* and *Neisseria meningitidis*^9,10^.

*Escherichia coli* has 8 DedA homologues (YqjA, YghB, YabI, YohD, DedA, YdjX, YdjZ, and YqaA), which are collectively essential^11^. Disruption of two of these genes, *yqjA* and *yghB*, results in a pleiotropic phenotype including altered lipid composition, cell division defects, sensitivity to elevated temperatures and alkaline pH, and susceptibility to multiple classes of antimicrobials^12–14^. Several observations have linked these phenotypes to defects in proton motive force maintenance^15,16^, suggesting a role for DedA proteins in membrane energetics. This interpretation is further supported by the sensitivity of a single Δ*yqjA* mutant to alkaline pH^13,14^. Furthermore, substitution of conserved acidic (Glu39/Asp51) or arginine (Arg130/Arg136) residues in YqjA render the protein unable to rescue the mutant phenotypes^17,18^, identifying these residues as functionally essential.

The link between the diverse membrane-associated phenotypes and DedA protein dysfunction has been illuminated somewhat by the discovery that several DedA family members possess lipid transport activity. In eukaryotes, DedA homologues TMEM41B and VMP1 have been shown to function as promiscuous lipid scramblases involved in the intra-organellar lipid trafficking^19,20^. In bacteria, the DedA homologue from *Bacillus subtilis* (PetA) has been proposed to transport phosphatidylethanolamine (PE) from the inner leaflet to the outer leaflet of the cytoplasmic membrane^21^. Furthermore, multiple studies have demonstrated that DedA homologues from *Vibrio cholerae*, *B. subtilis* and *Pseudomonas aeruginosa* can transport the 55-carbon isoprenoid lipid carrier undecaprenyl-phosphate (UndP) from the outer to the inner leaflet in the final precursor recycling step of peptidoglycan synthesis^22–25^. As UndP is required for Ara4N modification of Lipid A^26^, which is required for CAMP resistance, these findings provide a potential mechanistic link between DedA-mediated UndP transport and antimicrobial resistance^27–29^. Consequently, DedA proteins have emerged as attractive targets for the development of novel antimicrobial agents. However, progress towards this goal has been stymied by the limited structural and functional information available for this protein family.

To date, structural information for DedA proteins has been very limited. No experimentally determined DedA structures are currently available in the Protein Data Bank, and consequently structural insights have relied on computational models^30–32^. In earlier work, we validated the predicted topology of YqjA using substituted cysteine accessibility measurements (SCAM), providing the first experimental support for a structural model of a bacterial DedA protein^32^. Several observations suggest that DedA proteins may function as oligomers. Cysteine crosslinking studies of YqjA identified intermolecular interactions involving residues within or adjacent to the C-terminal transmembrane helix, while SDS-PAGE analysis revealed the presence of higher molecular weight species consistent with oligomer formation^32,33^. Further evidence for DedA oligomerisation and its functional role has emerged from studies of the DedA proteins UptA and PA4029, which appear to exist in a monomer-dimer equilibrium in solution^22,25^. For UptA, dimer formation correlates with enhanced UndP binding and is influenced by environmental pH, suggesting that oligomeric state may regulate DedA activity^22^. Collectively, these findings suggest that oligomerisation may be a conserved and functionally important feature of DedA proteins. However, the architecture of DedA oligomers and the relationship between oligomerisation, lipid binding and function remain unresolved.

Here, we use analytical ultracentrifugation, site-specific cysteine crosslinking, structural modelling and functional analyses to investigate the oligomeric state and architecture of YqjA. Our findings identify the most likely dimer arrangement of YqjA and reveal a putative lipid-binding site at the dimer interface, providing new insights into how DedA proteins may interact with membrane lipids.

## Results

### Molecular weight analysis reveals YqjA is predominantly a dimer

We previously published a structural model of monomeric YqjA, which was experimentally validated using site-specific cysteine labelling^32^. This modelling predicted that YqjA contains three transmembrane helices and two re-entrant hairpins (Fig.1a,b). To provide further insight into the architectural arrangement of YqjA, we first sought to determine its oligomeric state, which would provide fundamental structural information and key constraints for structural modelling. To determine the oligomeric state of YqjA, we overexpressed *yqjA* from an arabinose-inducible pBAD vector, in-frame with a C-terminal octa-histidine tag in *E. coli* C43 cells. We extracted YqjA with dodecyl-β-D-maltoside (DDM) detergent and purified the protein using IMAC followed by SEC (Fig. 1c). SDS-PAGE analysis of SEC-purified YqjA revealed the presence of a dominant protein band at ∼20 kDa (Fig. 1c, inset). The estimated molecular weight of YqjA based on the amino acid sequence is ∼26.3 kDa. However, integral membrane proteins often migrate faster in SDS-PAGE than expected^34^. Therefore, we reasoned that the band at ∼20 kDa represented monomeric YqjA. We also observed higher molecular weight bands at ∼40 and ∼60 kDa (Fig. 1c, inset), as seen previously^33^, which we hypothesised were dimeric and trimeric oligomers of YqjA, respectively. These data suggested that YqjA formed an oligomeric complex but provided no clues as to YqjA’s actual oligomeric state. To provide this information, we determined the sedimentation coefficient of the protein complex using sedimentation velocity analytical ultracentrifugation (SV-AUC), which has been applied to characterise numerous membrane proteins and transporters^35–38^. The sedimentation coefficient is influenced by several intrinsic properties of the particle, including its molecular mass, density, and shape.

**Figure 1.**
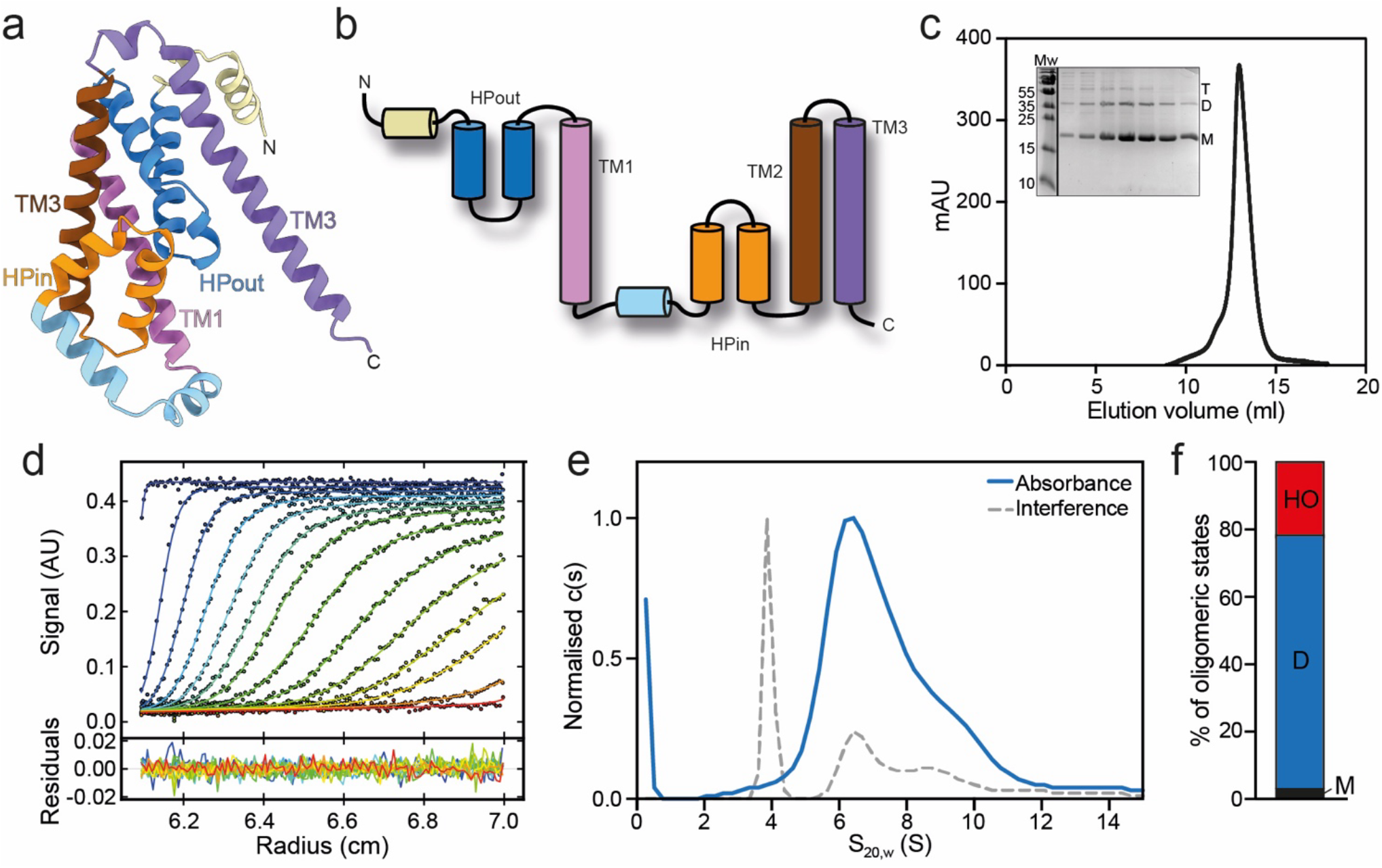
SV-AUC reveals YqjA is predominantly dimeric. **a)** Cartoon representation of a AF2 model of YqjA monomer. **b)** Topology map of YqjA monomer colour coded to match a). **c)** Size exclusion chromatography (SEC) A280 trace of IMAC-purified YqjA. *Inset*, SDS-PAGE analysis of the fractions taken across the SEC peak (11-15 ml elution volume). Molecular weight standards (Mw) are indicated, and molecular weights are shown in kDa. Presumed monomeric (M), dimeric (D) and trimeric (T) YqjA bands are labelled. **d)** Superposition of selected experimental data (dot) and fitted (line) sedimentation velocity profiles acquired in absorbance at 280 nm, during ≈ 12.5 hr at 130,000 rcf (top) and residual (bottom), for YqjA at 1.3 mg/ml. **e)** Normalised c(s) distributions obtained at 280 nm (continuous blue line) and using interference detection (dashed grey lines), for YqjA at 1.3 mg/ml. **f)** Percentages of total absorbance assigned to monomeric (M), dimeric (D) or higher oligomeric (HO) YqjA at 1.3 mg/ml. Higher oligomer fraction is a combination of absorbances consistent with potential 4-, 6-, 8- and 16-mers.

We analysed purified YqjA in DDM using SV-AUC at a concentration of 1.3 mg/ml, which revealed a predominant species sedimenting at S_20,w_ of ∼6.7 S in absorbance and in interference, which accounted for ∼76% of the total absorbance signal (Fig. 1d-f, SI Fig. 1c). Additionally, less abundant species were detected at higher sedimentation coefficients, suggesting the presence of larger assemblies. Moreover, we observed a peak, mainly in the interference optics, at S_20,w_ of ∼4 S (Fig. 1d-f, SI Fig. 1c). Hydrodynamic modelling, incorporating the experimentally estimated amount of bound detergent, identified the S_20,w_ ∼6.7 S as consistent with the dimeric protein with 2.5 g/g of bound DDM. The peak at S_20,w_ ∼4 S species is consistent with co-sedimenting free DDM micelles and low amounts of monomeric protein bound to detergent. AUC was also performed on more concentrated protein samples (2.6 mg/mL and 5.9 mg/mL), and analysis of the interference signal revealed that while the dimeric species remained the most abundant, the relative proportion of the peak at S_20,w_ of ∼4 S, corresponding to monomer, decreased (SI Fig. 1a-c). Taken together, these data strongly suggested that YqjA is predominantly dimeric in the detergent solution, and our data are consistent with YqjA existing in monomer-dimer equilibrium, as seen for another member of the DedA family^22,25^.

### Identification of candidate YqjA dimer interfaces using multiple AlphaFold approaches

Having established that YqjA is predominantly dimeric, we used multiple AlphaFold2 versions and AlphaFold3 to model the dimer interface and provide a structural framework for experimental validation. Together, these approaches generated six candidate models of the YqjA dimer interface (Fig. 2).

**Figure 2.**
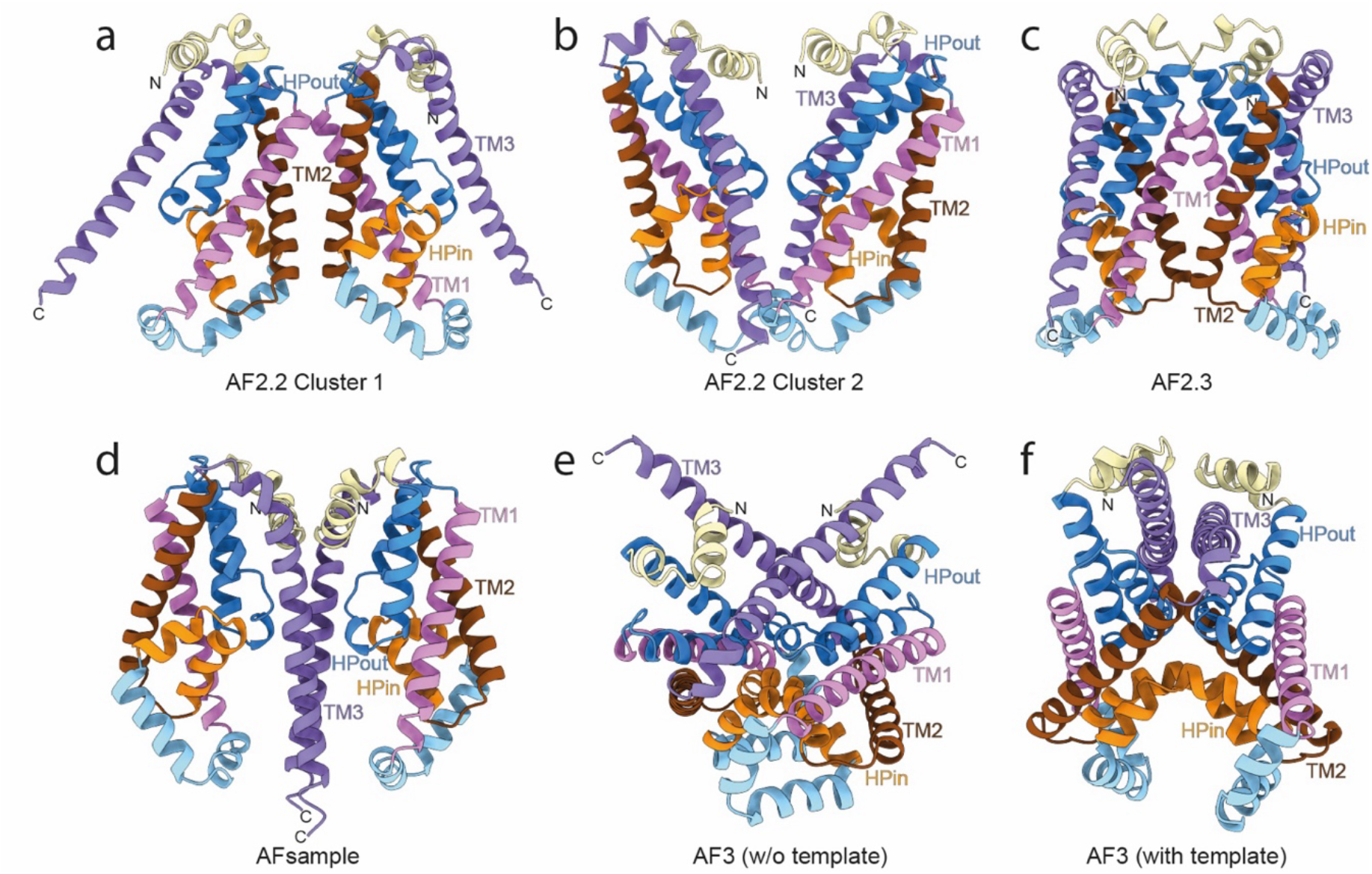
Candidate YqjA dimer models generated with AlphaFold variants. Cartoon representation of final structures selected from **a)** AlphaFold 2.2 cluster 1, **b)** AlphaFold 2.2 cluster 2, **c)** AlphaFold 2.3, **d)** AlphaFold sample, **e)** AlphaFold 3 without template and **f)** AlphaFold 3 with template. Models are coloured as in Fig 1a,b. N- and C-termini are indicated and transmembrane regions and re-entrant hairpins for the righthand protomer are labelled in each dimer.

Initially, we used AlphaFold v2.2 (AF2.2)^39,40^, which predicted two possible dimer interfaces, both with low confidence (Fig. 2a,b). Most models predicted with AF2.2 had very low combined pTM/ipTM scores with only a few above 0.2 (SI Fig 2a), indicating poor interface resolution (maximum ipTM value is 0.24), reflecting AF2.2’s inability to generate a good-quality model of the YqjA dimer. Indeed, only 19 out of 2500 predictions had AF combined ipTM+pTM score values of ≥0.2 (SI Fig 2a). Nevertheless, the protomers in each of these predictions have the same arrangement as the previous experimentally validated YqjA monomer model, providing some support for these models (SI Fig. 3a,b)^32^. The intersubunit interactions are predicted to be between the HPout re-entrant helices, TM1, TM2, and TM3 (SI Fig 4a). By measuring the minimum distance between these segments, we identified two clusters with two different dimer interfaces: one along TM2 of each protomer (cluster 1, Fig 2a) and the other along TM3 and the HPout re-entrant helices (cluster 2, Fig 2b). In the latter model, the protomers are proximal on the cytoplasmic side of the membrane but separate towards the periplasmic side, generating a V-shaped vestibule that provides direct access from the membrane bilayer to the tips of the re-entrant hairpins (Fig. 2b). However, according to the predicted local distance difference test (pLDDT), the confidence in the position of the helices at the dimer interface (SI Fig 5a,b) is better for the cluster 1 dimer (TM2/TM2’) than for the cluster 2 (TM3 & HPout/ TM3’ and HPout’).

Given the very low confidence of the dimers predicted using AF2.2 (mean 0.17 for best 20%), we modelled new structures with AlphaFold v2.3 (AF2.3), which consistently predicted a 3^rd^ possible dimer interface (Fig. 2c). Here, we obtained predictions that had, on average (mean 0.31 for the best 20%), higher scores than the AF2.2 predictions, but still below confident values (SI Fig. 2b, average for prediction of best neural network 0.29 for AF2.3 and 0.16 for AF2.2). Again, the protomer topology is very similar to the one predicted by AF2.2 and previous models (SI Fig. 3c)^32^. The contact map of the score-filtered predictions (310 out of 500) reveals consistent interactions between TM1 and TM2 (SI Fig. 4b). Clustering based on the minimum distance between interacting helices shows one predominant interface (one cluster represents ∼70% of all filtered structures) along TM1 and HPout regions (Fig. 2c) that, according to pLDDT score, are reasonably predicted (SI Fig. 5c).

Since neither of the two versions of AlphaFold 2 could confidently predict YqjA dimers, we used a third AlphaFold-based approach that allowed us to modify neural network parameters to sample other variables not accessible with the standard AlphaFold algorithms. We performed six independent AFsample (based on AlphaFold 2) modelling runs with different specific options (see Methods and SI Table 1 for details), testing score convergence of each separate set (SI Fig. 2c-h). AFsample predicted a fourth alternative dimer interface (Fig. 2d), which had a higher score than any of the previous AlphaFold 2 predictions (SI Fig. 2c-h). We retained a reasonable number of predictions after filtering by combining pTM+ipTM score (419 out of 800). In this case, the most populated cluster (93% of the filtered structures) showed extensive contacts between the TM3 of each protomer (SI Fig. 4c). Although the per-residue confidence of this region is low (SI Fig. 5d). Interestingly, this model contains a large cavity between the protomers on the cytoplasmic side of the membrane, whereas the periplasmic side of the interface is largely occluded (Fig. 2d).

**Table 1.**
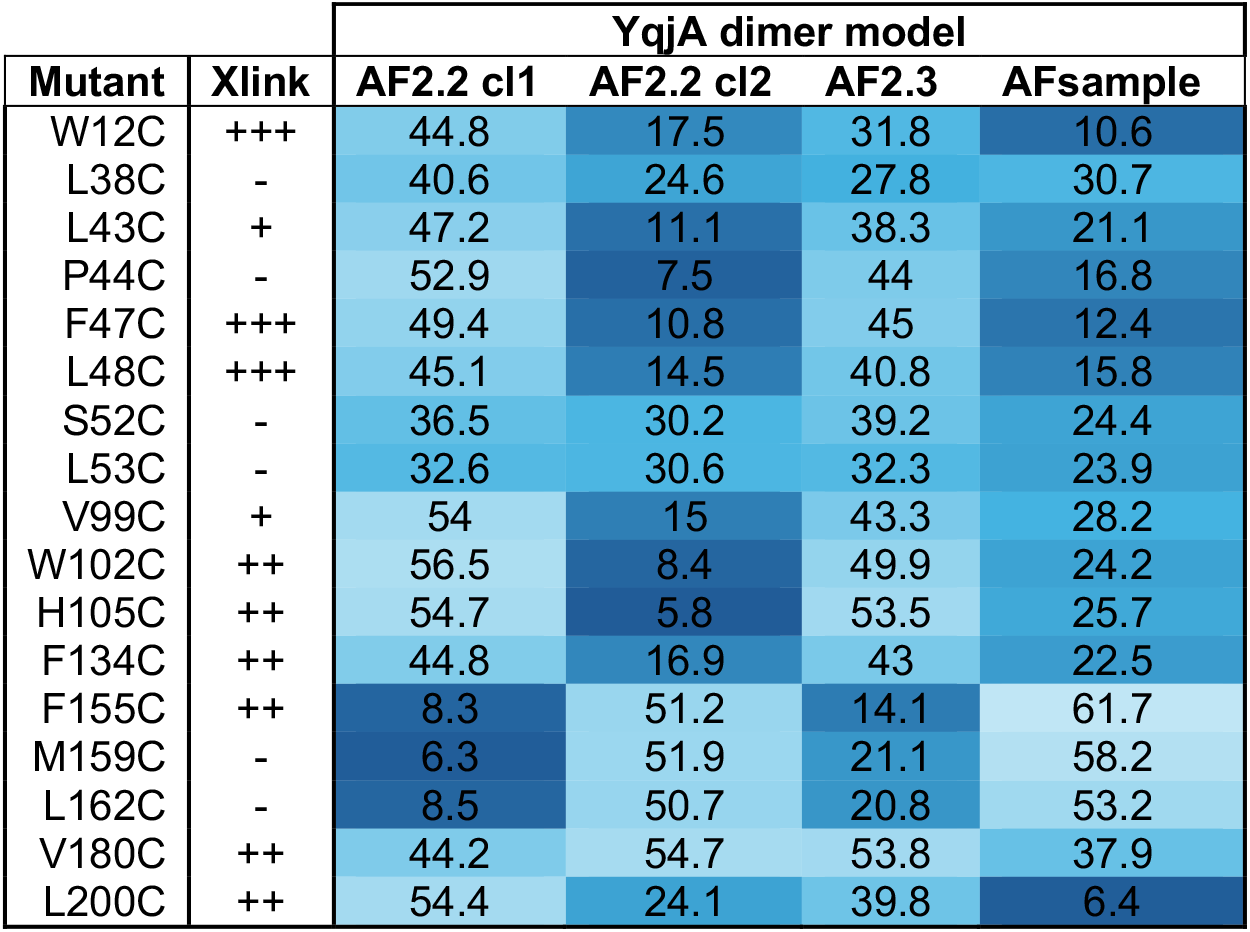
Intermolecular Cα–Cα distances and disulfide crosslinking efficiencies of YqjA cysteine-substitution mutants. Distances are shown in Å. Crosslinking efficiency is indicated as follows: –, none; +, weak; ++, moderate; +++, strong. Distance values are colour-coded from dark blue (shortest distance) to light blue (greatest distance).

Although AFsample predicted better-scoring models, the score values were still modest (best values ∼0.4). At this time, AlphaFold 3 became available; therefore, we expanded our YqjA dimer predictions using AlphaFold 3. The four templates found by AlphaFold (4ZV6, 1IIJ, 2L9U, 6UUR) were highly unrelated to each other (de novo protein, NMR solution of portion of transmembrane proteins, and prion protein); we tested the effect of templates by performing two separate runs: one with structural templates and one without. These runs yielded the highest-scoring models of the entire set (SI Fig. 2i,j), and according to the pLDDT scores, were of reasonable confidence (SI Fig. 5e,f). However, unlike the AF2.2, AF2.3 and AFsample models, whose intermolecular contact hotspots were well-defined (SI Fig. 4a-c), the contacts in the AF3 models were spread across much of the opposing protomer surfaces (SI Fig. 4b), resulting in a poorly defined dimer interface (Fig. 2e,f). In addition, models of both AF3 approaches were largely inconsistent with our previous SCAM data. Most notably, these AF3 models predicted the N-and C-termini to reside on the same side of the membrane, whereas our experimental data place them on opposite sides (SI Fig. 3e,f)^32^. Based on these observations, neither of these AF3 models was considered further as a plausible representation of the YqjA dimer.

Given that the modelling strategy generated multiple possibilities and did not converge into a single predominant interface, we used cysteine crosslinking to determine which of the structural models was the most accurate depiction of the YqjA dimer.

### Mapping of YqjA’s dimer interface using site-specific cysteine crosslinking

To determine which structural model most accurately represented the YqjA dimer, we identified interface residues whose relative positions differed between the predicted models. Analysis of the C*a*-C*a* distances between the equivalent residues in the two protomers for each of the structural models revealed substantial variation, with some positions exhibiting separations ranging from ∼6 to 50 Å, depending on the model. This analysis identified a series of positions at which cysteine substitution and crosslinking could be used to discriminate between the proposed dimer arrangements. Based on these criteria, 17 residues were individually substituted with cysteine in the cysteine-free version of YqjA (YqjA_cysless_) that we previously showed was functionally active (Table 1)^32^. As the dimer interface predictions were generally of low confidence, we adopted relatively permissive distance criteria when selecting candidate residues, including some positions whose predicted C*a*-C*a* separations exceeded those commonly observed for disulfide-bonded cysteine pairs (typically 5-7 Å)^41^. Upon oxidation with copper phenanthroline (CuPhen), sufficiently proximal cysteine pairs would form an intermolecular disulfide that could be detected as a band shift in SDS-PAGE analysis.

To ensure any structural information derived from our crosslinking study was physiologically meaningful, we first determined whether the cysteine mutants were properly folded and functional. To do this, we expressed the mutated genes from the pBAD expression vector and assessed their ability to restore growth to a temperature-sensitive *E. coli* W3110 mutant devoid of the *yqjA* and *yghB* genes (W3110ΔΔ), which grows poorly at 42°C^42^. While expression of a negative control gene (aspartate transporter, Glt_Ph_) could not restore growth at the non-permissive temperature, YqjA_wildtype_, YqjA_cysless_, and all the cysteine mutants restored growth demonstrating that they were functionally active and folded (SI Fig. 6).

To perform the cysteine crosslinking, we individually expressed the members of the mutant library in *E. coli*, which were either left untreated, reduced using dithiothreitol (DTT) or oxidised by addition of CuPhen, which would promote disulfide crosslinking. Following these treatments, all samples were treated with an excess of methyl methanethiosulfonate (MMTS) prior to SDS-PAGE, which would alkylate any unmodified cysteines to prevent aberrant in-gel dimer formation between SDS-unfolded proteins. We monitored crosslink formation using SDS-PAGE and Western blotting with an anti-his antibody. Analysis of the Western blots for each mutant revealed stable dimer formation for 9 of the mutants; W12C, F47C and L48C exhibited high levels of crosslinking upon addition of CuPhen, whereas more moderate crosslink formation was observed for W102C, H105C, F134C, F155C, V180C and L200C under the same conditions (Fig. 3a). The remaining cysteine mutants either exhibited very low levels of crosslink formation, as is the case for V99C and L43C, or no crosslinking formation at all was observed. From this, we concluded that the residues exhibiting high or moderate levels of crosslinking identify regions of YqjA that are likely brought into close proximity upon dimer formation and provide a set of experimental constraints for evaluating the predicted dimer models.

**Figure 3.**
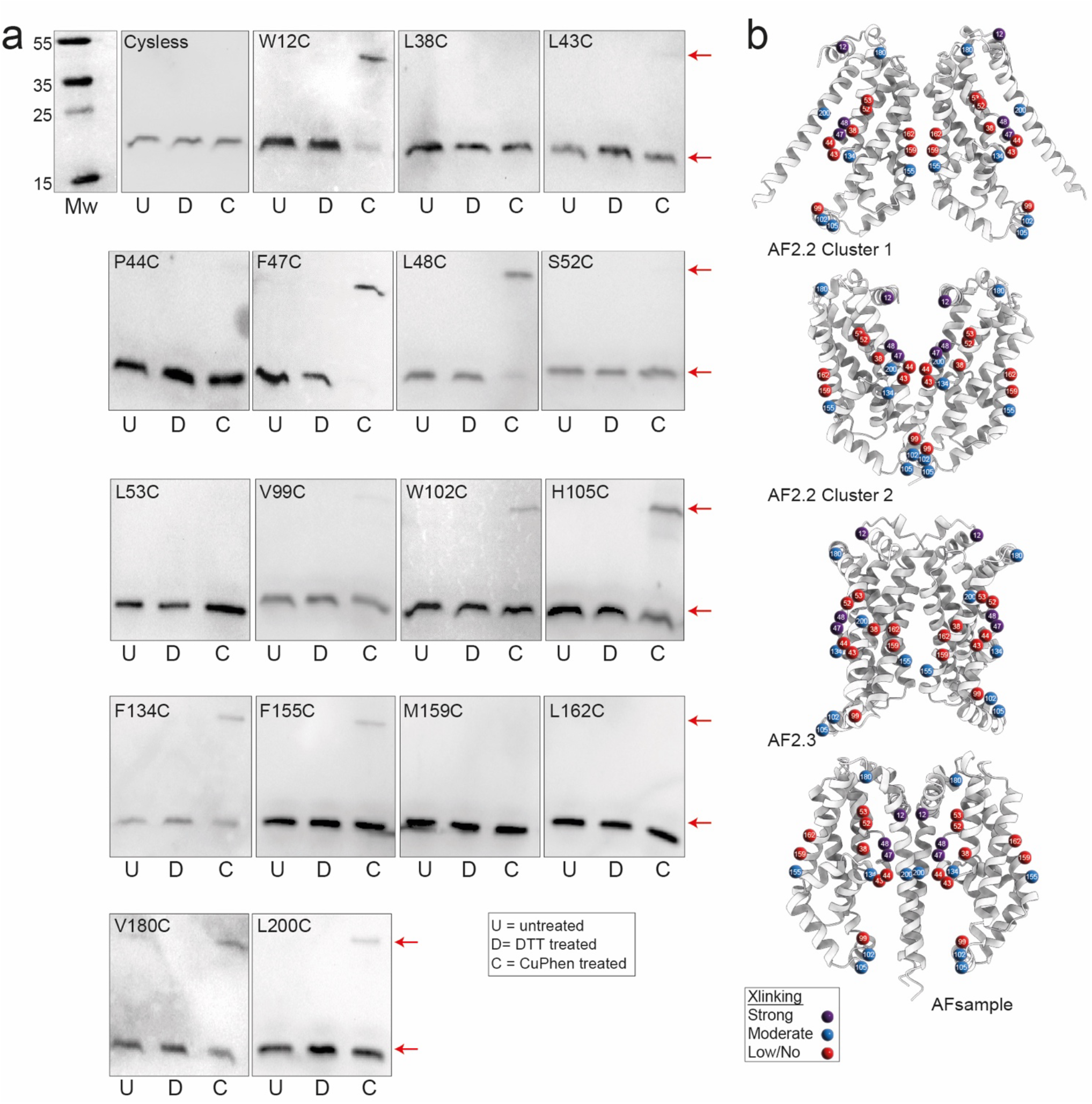
Disulfide crosslinking of YqjA and mapping of crosslinking residues onto structural models. **a)** Western blot analysis of YqjA cysteine-substitution mutants following no treatment (U), treatment with 5 mM DTT (D) or treatment with 0.6 mM CuPhen (C). Red arrows indicate the monomer and dimer YqjA bands. Molecular weight standards (Mw) are indicated, and molecular weights are shown in kDa. Representative blots from three independent biological replicates are shown. **b)** Mapping of crosslinking residues onto the four predicted YqjA dimer models. Residues exhibiting strong (purple spheres), moderate (blue spheres), or low/no crosslinking (red spheres) in the CuPhen assays are indicated. Crosslinking efficiency was assigned based on the relative abundance of dimer observed in the Western blot analyses shown in panel a).

### Crosslinking analysis suggests conformational diversity within the YqjA dimer interface

To determine which of the predicted models was most consistent with the CuPhen crosslinking data, we mapped the residues exhibiting high or moderate levels of crosslinking onto each model (Fig 3b, Table 1). Firstly, the positions and intermolecular distances of the strongly crosslinking residues (W12, F47 and L48) indicate that the AF2.2 cluster 1 and AF2.3 models are incompatible with the experimental data. In addition, of the residues exhibiting moderate crosslinking, only F155 contributed to the dimer interface in either model (Fig 3b, Table 1). Taken together, the poor agreement between these models and the crosslinking data leads us to discount them from further consideration.

Having excluded AF2.2 cluster 1 and AF2.3 models, our analysis narrowed to AF2.2 cluster 2 and AFsample models, both of which were in generally good agreement with the crosslinking data, placing all three strongly crosslinking residues at the dimer interface (Fig 3b). In both the AF2.2 cluster 2 and AFsample models, F47 and L48, were positioned at the centre of the dimer interface, where they formed part of the re-entrant hairpins (HPout) contributed by each protomer (Fig. 3b). Similarly, F134, which is located within the cytoplasmic re-entrant hairpin (HPin), occupied a comparable position in both models, lying between the two protomers immediately beneath F47 and L48. While the exact positioning of these residues differed somewhat between AF2.2 cluster 2 and AFsample models, their overall arrangement within the central portion of the dimer interface was conserved. The principal differences between the two models occurred outside this central hairpin region. In the AFsample model, W12C residues within the N-terminal periplasmic helices were positioned in much closer proximity than in AF2.2 cluster 2 (Fig. 3b). In contrast, residues W102 and H105 on the cytoplasmic face showed the opposite trend, being closer together in AF2.2 cluster 2 and more widely separated in AFsample. Thus, while the arrangement of the re-entrant hairpins was largely conserved, the relative positioning of the N-terminal and cytoplasmic regions differed substantially between the two models.

Three residues represented exceptions to the overall trend. F134C exhibited moderate crosslinking despite relatively large predicted separations in both AF2.2 cluster 2 and AFsample (16.9 and 22.5 Å, respectively; Table 1). Similarly, F155C exhibited moderate crosslinking despite being positioned near the dimer interface only in models that were otherwise thoroughly inconsistent with the experimental data, whereas V180C exhibited moderate crosslinking despite remaining relatively distant in all models examined (∼38-54 Å Fig. 3b, Table 1). We reasoned that these observations may reflect limitations of the static structural models, the formation of low-frequency intermolecular contacts not captured by the predicted structures, or the presence of transient conformations arising from the monomer– dimer equilibrium exhibited by YqjA.

Taken together, these data support a dimeric architecture for YqjA in which the re-entrant hairpins form the core of the dimer interface. Although AF2.2 cluster 2 and AFsample differ in the relative positioning of several peripheral helices, both models place the conserved acidic (E39/D51) and arginine (R130/R136) residues at the centre of the assembly. Given the established role of several DedA proteins in lipid transport and the central location of the conserved acidic and arginine residues within the YqjA dimer, we investigated whether these residues might contribute to lipid binding.

#### Lipid-bound models reveal a putative phospholipid-binding site within the YqjA dimer

As lipid binding has been proposed for other DedA family members^21–25,43^, we used AF3 to generate lipid-bound models of YqjA dimer. To do this, we co-folded two molecules of YqjA with a molecule of 1-palmitoyl-2-oleoyl-sn-glycero-3-phosphoethanolamine (POPE), a representative phosphatidylethanolamine (PE), or 1-palmitoyl-2-oleoyl-sn-glycero-3-phospho-(1’-rac-glycerol) (POPG), representing the phosphatidylglycerol (PG) family, the two most abundant lipid classes in the *E. coli* lipidome.

These predictions yielded relatively well-scored models (Fig. 4a, SI Fig 7a,e), allowing us to filter more stringently than with any of the AlphaFold2 variants; pTM/ipTM scores above 0.4 were obtained for 670 and 153 models per 1000 predictions for the POPE- and POPG-bound dimers, respectively (SI Fig. 7a). Importantly, the majority of filtered models adopted a parallel arrangement (SI Fig 7b); ∼80% and 90% of predictions for the POPE- and POPG-bound models, respectively, and were highly consistent with our previous SCAM data (SI Fig. 7c)^32^. Interestingly, inclusion of POPE or POPG during AF3 modelling led to a remarkably different intermolecular contact pattern prediction (SI Fig. 7d). Rather than the diffuse contacts observed in the lipid-free AF3 models (SI Fig. 4d,e), the lipid-bound models exhibited a much more localised interaction interface characterised by discrete interaction hotspots (SI Fig. 7d).

**Figure 4.**
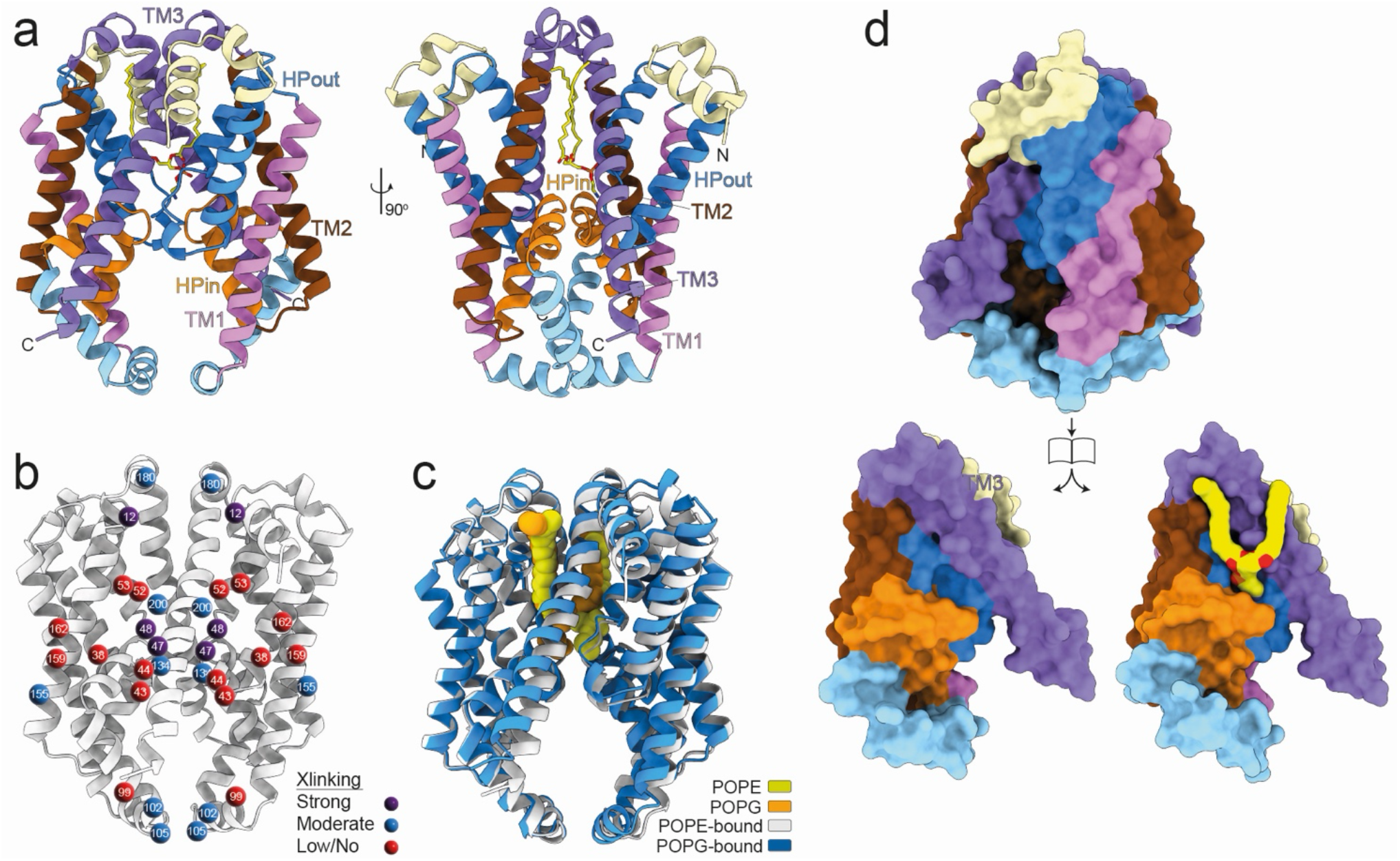
Lipid-bound AF3 models. **a)** Cartoon representation of the POPE-bound AF3 model of the YqjA dimer. The POPE molecule is represented as yellow sticks. Two views of the model are shown and the protein is coloured as in Fig. 2. **b)** Mapping of crosslinking residues onto the POPE-bound AF3 YqjA dimer model. Spheres represent tested cysteines and are coloured as in Fig. 3b. **c)** Superposition of cartoon representation of POPE- (white) and POPG-bound (blue) AF3 models. POPE (yellow) and POPG (orange) are represented as spheres. **d)** Surface representation of the YqjA dimer:POPE complex (top) and the dimer interface of both protomer if opens as a book. POPE is represented as yellow spheres.

The contact maps show a persistent interaction between TM3 and HPin of both protomers (SI Fig 7d). Clustering based on the minimum distance between these interacting regions reveals the same preferred interface and lipid-binding site in models with both lipid types. The most populated cluster contains 144 models for POPE-bound YqjA and 37 for POPG-bound YqjA. The dimerisation interface and the protein regions contacting the lipid were predicted with high confidence in the backbone prediction (defined as pLDDT above 70% across most of this region, SI Fig 7e). This suggests that, even though we may lack the resolution to unambiguously define specific protein-lipid interactions, the overall arrangement can be predicted with confidence.

Consistent with the AF2.2 cluster 2 and AFsample models obtained without lipid, all strongly and most moderately crosslinking residues were located at the dimer interface in the lipid-bound models (Fig. 4b). Agreement with the experimental crosslinking data was also improved for F134C and V180C. Most notably, F134C, which exhibited moderate crosslinking experimentally, was positioned >16.9 Å from the opposing protomer in all lipid-free models but only 7.3-7.7 Å apart in the lipid-bound models. Although the predicted separation of V180C remained relatively large (18-19 Å), this represented a substantial improvement over the >37 Å predicted by all lipid-free models. The predicted structures of the POPE and POPG-bound models are essentially identical (RMSD of 0.6 Å across all 220 residues, Fig. 4c). Consistent with the dramatic difference in the contact map when co-folded with lipid (SI Figs. 4d,e and 7d), the lipid-bound AF3 models adopted a considerably different dimer arrangement compared to the initial AF3 models (Fig. 4a). Structural comparison revealed that the lipid-bound models most closely resemble the AF2.2 cluster 2 architecture, with the principal difference arising from a rigid-body movement of one protomer relative to the other that occludes the periplasmic vestibule (Fig. 4a). Indeed, superposition of the POPE-bound model protomer with protomers from the AF2.2 cluster 2 and AFsample models revealed RMSDs of 0.8 Å across 153 and 141 residues, respectively (SI Fig. 8a). The primary difference in the protomers of the three protomers is the positioning of the C-terminal TM3, which adopts distinct orientations in each prediction (SI Fig. 8a), as if pivoting around its N-terminal end, thus exposing different parts of the binding site.

Both lipid-bound models predict that the lipid molecule binds at the interface between the two protomers, with their aliphatic chains sandwiched between the TM3 of the two protomers and extending towards the periplasmic side of the membrane (Fig. 4a,c,d). The lipid headgroups are positioned adjacent to the re-entrant hairpin loops where they interact with the tips of HPin and HPout (residues 39-51 and 132-135, respectively, Fig. 4a,c, 5a,b). The interaction is asymmetric, with the lipid headgroup displaced towards one side of the dimer interface (Fig 4c,d). Interestingly, structurally aligning the POPG- and POPE-bound models revealed that the lipid headgroups are not predicted to interact with the same protomer (Fig. 4c), suggesting that lipid binding can be coordinated by residues contributed from both subunits of the dimer.

In our POPE-bound model, the side chains of Glu39 and Asp51 are positioned to form potential salt-bridge interactions with Arg136 (3.16 Å and 2.40-3.13 Å, respectively), while Arg136 itself lies sufficiently close to the POPE phosphate group (3.81 Å) that a modest local rearrangement could permit direct interaction (Fig. 5a). Together, these functionally important residues form a putative interaction network adjacent to the lipid headgroup. The only clear direct interaction with POPE in the static model is provided by Ser52, which is positioned to hydrogen bond with a phosphate oxygen (2.92 Å). In contrast, Arg130 points away from the binding site and has no apparent role in lipid coordination (Fig. 5a). The predicted headgroup interactions differ somewhat in the POPG-bound model (Fig. 5b). As in the POPE-bound model, Ser52 is positioned to hydrogen bond with a phosphate oxygen of POPG (3.3 Å distance, Fig. 5b). However, here, Glu39 directly contacts the glycerol moiety of the POPG headgroup through a predicted hydrogen bond between its sidechain carboxylate and a glycerol hydroxyl (2.69 Å, Fig. 5b). Although Arg136 does not directly contact the POPG headgroup, its guanidinium group lies within 3 Å of the carboxylate groups of Glu39 and Asp51, suggesting it is likely the hub of a salt bridge network (Fig. 5b).

**Figure 5.**
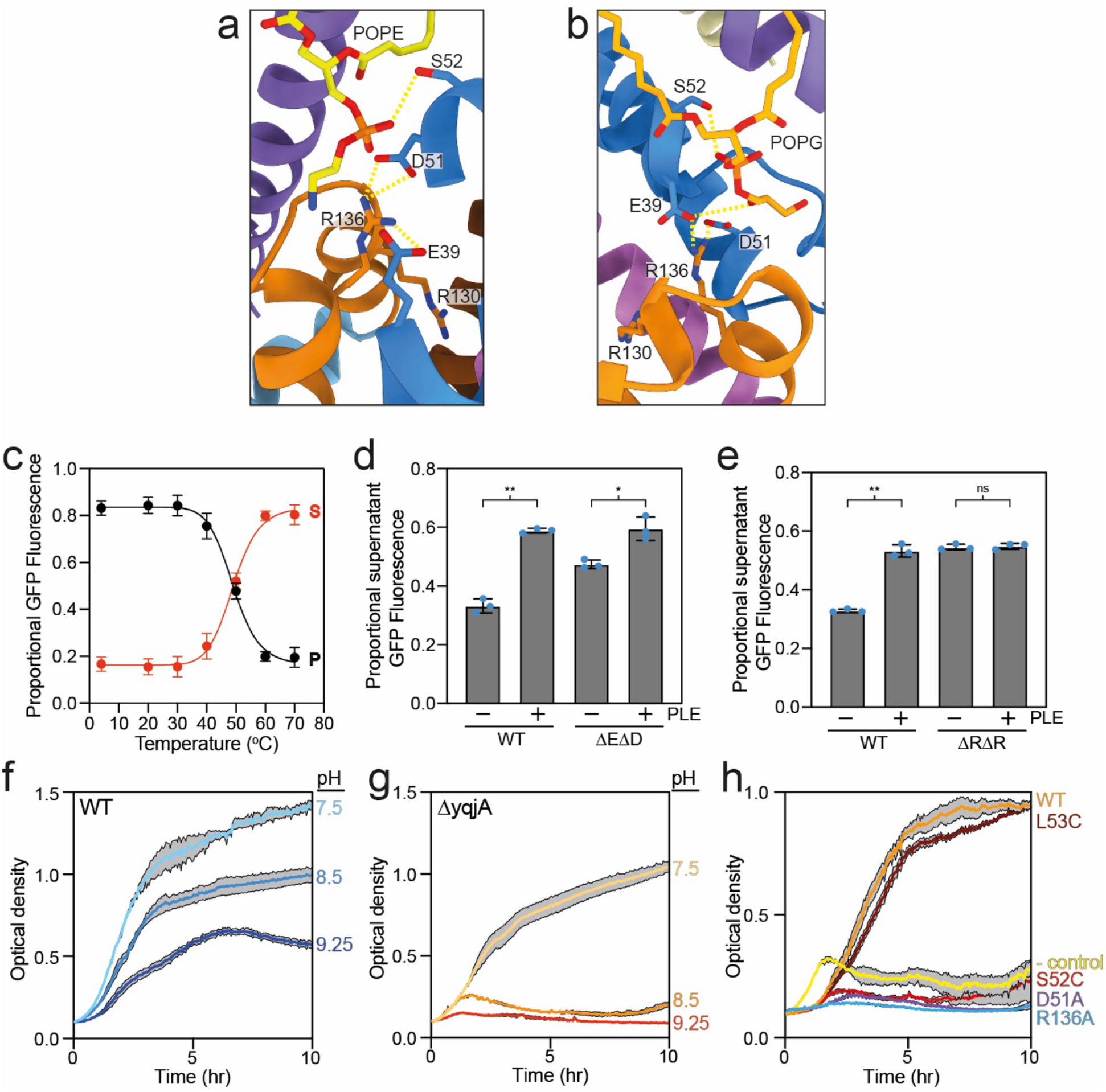
Experimental validation of the predicted lipid-binding site. a-b) Close-up view of the predicted lipid headgroup binding sites for the POPE- **(a)** and POPG-bound **(b)** models. Structures are coloured as in Fig 4a, predicted interactions are indicated (dashed yellow lines), and relevant residues and lipids are labelled. **c)** Proportion of GFP fluorescence in the pellet (black data) and supernatant (red data) as a function of incubation temperature for wildtype YqjA. Data plotted is the mean of 9 replicates and error bars represent SD. **d-e)** Proportion of the GFP fluorescence remaining in the supernatant after heating at Tm + 5°C for YqjA-WT, ΔEΔD mutant in 50 mM NaCl **(d)**, and YqjA-WT and ΔRΔR mutant in 25 mM NaCl **(e)**. Bars represent the mean of three independent experiments, each performed in technical triplicate, error bars represent SD, and single data points are shown in blue. Statistical significance was assessed using a two-tailed unpaired Welch’s t-test. \**P* < 0.05, \*\**P* < 0.01; ns, not significant. **f-g)** Growth of *E. coli* BW25113 wildtype **(f)** and *E. coliΔyqjA* **(g)** at 37°C in LB set to pH, 7.5, 8.5 and 9.25. **h)** Growth of *E. coliΔyqjA* harbouring plasmids expressing *yqjA* wildtype (WT, orange), L53C mutant (maroon), S52C (red), D51A (purple), R136A (blue) or control plasmid expressing an unrelated gene (yellow). For f-h, the average of triplicate data set is shown, error is represented by grey shading and represented SD.

To support this model, we sought to assess whether YqjA was able to bind lipids. To do this, we used a lipid-compatible thermal shift assay, GFP-TS (GFP thermal stability assay)^44^, which has been used previously to monitor substrate and lipid interactions with multiple transport proteins^45,46^. This approach entails heating a GFP-tagged protein variant in the presence of the detergent, n-Octyl-β-D-Glucopyranoside (β-OG), which promotes aggregation of unfolded protein^44,45^. The heat-denatured aggregates are sedimented by ultracentrifugation, and the partitioning of GFP fluorescence between the supernatant and pellet reflects the extent of protein denaturation at each temperature, from which an apparent T_m_ can be determined.

To enable use of this approach, we fused GFP to the C-terminus of YqjA and confirmed that the GFP-tagged protein remained fully functional using a phenotype rescue assay (SI Fig. 9a). Using this construct, we determined the apparent T_m_ of wildtype YqjA (YqjA-WT) in membranes using GFP-TS by quantifying the distribution of GFP fluorescence between the supernatant and pellet following incubation at a range of temperatures (Fig. 5c). This analysis revealed an apparent T_m_ of 49 ± 1°C for YqjA-WT (Fig. 5c). We next sought to investigate whether the addition of lipids could stabilise YqjA-WT, which would indicate protein:lipid interactions. We compared the thermostability of YqjA-WT in the presence and absence of additional lipid by quantifying the fraction of protein that resisted aggregation after incubation at T_m_ + 5°C. As the fraction of protein remaining soluble at T_m_ + 5°C reflects relative protein stability, this approach allows comparison of multiple conditions without generating full melting curves. As we do not yet know the lipid specificity of YqjA, we opted to test lipid stabilisation effects using *E. coli* polar lipid extract (EPL), which is composed of: 67% PE, 23.2% PG, 9.8% cardiolipin (according to the manufacturer’s website). Addition of 5 mg/ml EPL to solubilised YqjA-WT-containing membranes led to a significant increase in the amount of YqjA-WT in the supernatant from 0.3 to 0.6 of the proportional fluorescence, indicating substantial stabilisation (Fig. 5d). Interpreted at face value, this result suggests that YqjA does indeed interact with lipids. However, because YqjA is an integral membrane protein, the observed stabilisation may arise from a general stabilising effect of EPL on the membrane protein-detergent complex rather than from a specific interaction with YqjA. To address this, we introduced mutations into the YqjA-GFP construct that were predicted to disrupt the lipid-binding site and assessed whether the stabilising effect was retained. To this end, we generated two double mutants: E39A combined with D51A (ΔEΔD) and R130A combined with R136A (ΔRΔR), both of which should be non-functional variants based on phenotypic analysis^18,33^. The GFP-TS melt curves for the ΔEΔD mutant revealed a T_m_ of 58.7 ± 0.6°C for (SI Fig. 9b), ∼9.7°C higher than wildtype. For the ΔRΔR mutant, under our standard conditions, we initially obtained Tm values over 70°C, which we considered unreliable due to the proximity to the T_m_ for GFP (76°C)^47^. To bring the ΔRΔR Tm within a reliable range, we reduced the NaCl concentration from 50 to 25 mM, which revealed apparent T_m_s of 49.75 ± 0.3°C for wildtype and 58 ± 5.6°C for ΔRΔR (SI fig. 9c,d). To assess the impact of additional lipid on the thermostability of the double mutants, we incubated them at T_m_ + 5°C in the presence and absence of EPL. These data revealed that, although addition of EPL stabilized ΔEΔD, the effect was substantially smaller than that observed for YqjA-WT (Fig. 5d). More starkly, addition of EPL to ΔRΔR had no effect on the apparent protein stability (Fig. 5e), suggesting that these mutations disrupt the interaction responsible for lipid-mediated stabilisation. Together, these data provide additional support for the predicted structure of dimeric YqjA-lipid complex and suggest that YqjA specifically interacts with lipids via these conserved residues.

### Identification of a functionally important serine residue

Our analysis of the predicted lipid-binding site revealed a potential hydrogen bond between the side chain of S52 and the POPE/POPG headgroups (Fig. 5a,b), suggesting that this residue may play a functionally important role. However, the YqjA-S52C-producing plasmid rescued the temperature-sensitive phenotype of an *E. coli* strain lacking both *yqjA* and *yghB* (SI Fig. 6), indicating that the substitution does not abolish YqjA function. We therefore sought to assess the functional consequences of the S52 mutation in greater detail by determining whether it could also rescue the Δ*yqjA*-specific alkaline intolerance phenotype^14,48^. As has been previously observed^48^, wildtype *E. coli* exhibits reduced but substantial growth in media with elevated pH (Fig. 5f), whereas *E. coliΔyqjA* fails to grow at pH 8.5 or higher (Fig. 5g). To test whether S52 is involved in the function underlying alkaline tolerance, we transformed an *E. coli*Δ*yqjA* strain with plasmids encoding YqjA-WT, a non-transporter control (-control), YqjA-R136A, YqjA-D51A or YqjA-S52C. The resulting strains were grown in liquid medium adjusted to pH 8.5 (Fig. 5h). Consistent with previous literature^18^/09/2026 22:28:00, YqjA-WT supported robust growth at pH 8.5, whereas YqjA-D51A did not, nor did YqjA-R136A, indicative of their important functional role. Interestingly, Δ*yqjA* cells producing YqjA-S52C also failed to grow at pH 8.5, indicating a previously unidentified functional role for this residue. To investigate if this was a specific effect of mutating the Ser52 sidechain or general disruption of that region of the protein, we also assessed whether mutating the neighbouring residues L53, would affect growth at pH 8.5. Our growth assays reveal that this strain grow similarly to YqjA-WT producing cells, supporting the functional relevance of Ser52.

## Discussion

In this study, we demonstrate that the *E. coli* DedA protein YqjA exists predominantly as a dimer in solution. By combining analytical ultracentrifugation, site-specific cysteine crosslinking and structural modelling, we identified the most likely dimer architecture and showed that the conserved functionally important acidic and arginine residues are positioned at the centre of the dimer interface. Modelling of dimeric YqjA in the presence of phospholipids predicts a lipid-binding site located adjacent to these residues, while thermal stability and mutagenesis data provide independent support for lipid binding. Together, these findings provide new structural and functional insights into the DedA superfamily and support a role for lipid binding in YqjA function.

### YqjA is predominantly a dimer

Previous studies have suggested that DedA proteins can exist as dimers. Native mass spectrometry studies on the UndP-binding DedAs UptA from *B. subtilis* and PA4029 from *P. aeruginosa* indicated both proteins exist in a monomer-dimer equilibrium, with the dimeric formation favouring UndP binding^22,25^. Similarly, previous crosslinking studies of YqjA have identified intermolecular interactions involving residues in and around the C-terminal transmembrane helix, hinting that YqjA also oligomerises^32,33^. Here, using sedimentation velocity AUC, we have demonstrated that YqjA is predominantly dimeric in solution, although small populations of monomeric and higher-order oligomeric species were also observed (Fig. 1d-f, SI Fig. 1a-c). The presence of both monomeric and dimeric species is consistent with the monomer-dimer equilibria reported for UptA and PA4029^22,25^, suggesting that dynamic oligomerisation may be a conserved feature of DedA proteins. While the functional significance of this equilibrium remains unclear, the observation that dimer formation correlates with substrate binding in other DedA proteins raises the possibility that oligomerisation contributes directly to DedA function. Interestingly, oligomerisation of UptA was shown to be pH-dependent using native mass spectrometry, with dimer formation favoured at physiological and alkaline pH and reduced under more acidic conditions^22^. Whether YqjA exhibits a similar pH-dependent equilibrium remains unknown, but such behaviour could provide a mechanism for regulating DedA activity in response to changing environmental conditions and may help explain the alkaline-sensitive phenotypes associated with DedA dysfunction^14,48^.

### The conserved re-entrant hairpins define the functional core of a dynamic YqjA dimer

In the absence of experimentally determined DedA structures, computational models provide an important framework for interpreting DedA function. However, our results demonstrate the importance of experimental validation when evaluating these models. Although the AF3-derived models were assigned the highest confidence scores (SI Fig. 2 and 5), they were largely inconsistent with our previous SCAM analysis (SI Fig. 3), exhibited diffuse contacts across the protomer surface (SI Fig. 4), and produced a poorly defined dimer interface, revealing that model confidence alone was insufficient to identify the native dimer architecture. In contrast, the lower-confidence AF2.2 cluster 2 and AFsample models showed substantially better agreement with both the crosslinking data presented here and our previous SCAM analysis (Fig. 3, SI Fig. 3). Remarkably, co-folding the YqjA dimer with phospholipid using AF3 produced a strikingly different dimer architecture that converged on the same core interface identified by the AF2.2 cluster 2 and AFsample models (Fig. 4, SI Fig 7). Although the lipid-bound models did not fully reconcile all the experimental constraints, they retained the central re-entrant hairpin interface and further improved agreement with the crosslinking data for several residues, most notably F134C. While none of the three models fully explained every crosslinking observation, they all agreed that the re-entrant hairpins HPout and HPin formed the core of the dimer interface.

Despite the differences in dimer architecture, the individual protomers of the AF2.2 cluster 2, AFsample and lipid-bound AF3 models were remarkably similar. Superposition revealed RMSDs of <0.8 Å over more than 140 residues. In addition to some variation in the C-terminal helix position, the major differences between the three models arose from the relative positioning of the two protomers. The models can largely be interconverted by rigid-body movement of one protomer relative to the other, resulting in reciprocal opening and closing of the periplasmic and cytoplasmic vestibules while preserving the central re-entrant hairpin interface. We therefore tentatively hypothesise that these models do not represent alternative dimer architectures, but rather distinct conformational states of a common dimeric assembly. This interpretation is supported by our previous SCAM analysis, in which the interfacial residue S52 exhibited only partial protection from the membrane-impermeant reagent MTSES, suggesting that this region samples multiple conformations with different solvent accessibility^32^. Such reciprocal rearrangements are reminiscent of alternating-access transport mechanisms, in which substrate-binding sites are exposed sequentially to opposite sides of the membrane^49^. Interestingly, this proposed mechanism bears resemblance to the “trap-and-flip” mechanism described for the lipid flippases LtaA and MFSD2A^50,51,52^, in which lipids are sequestered from one leaflet, enclosing them in a central cavity and delivering the lipid to the opposing leaflet without headgroup tail being exposed to the core of the bilayer. However, experimental structures of DedA proteins will be essential to establish whether these predicted arrangements represent genuine intermediates within the YqjA transport cycle.

### A putative lipid-binding site at the YqjA interface

Our lipid bound models predict a phospholipid binding site at the YqjA dimer interface, positioned adjacent to highly conserved residues previously shown to be important for YqjA function. In both POPE and POPG-bound models, the lipid headgroup is proximal to the re-entrant hairpin loop and is surrounded by a network of conserved polar and charged residues, including Glu39, Asp51, and Arg136 (Fig. 5a,b). Although the precise interactions differ between the POPE- and POPG-bound models, Glu39, Asp51 and Arg136 form a potential interaction network adjacent to the lipid headgroup that could contribute to lipid recognition and coordination. Our GFP-TS data showing reduced lipid-induced protein stabilisation when these residues are mutated support their involvement in lipid binding (Fig. 5c-e). In addition, the importance of these residues to lipid binding has been identified in work on UptA from *B. subtilis*^22^, which demonstrated using native mass spectrometry that substitution of residues equivalent to Glu39, Arg130 and Arg136 (Glu32, Arg112 and Arg118 in BsUptA) substantially reduced UndP binding^22^. Combined, these data strongly suggest a role for this interaction network in lipid binding. Based on the linkage between YqjA and PMF maintenance and the presence of protonatable residues in the binding site, it is possible that one or both of the acidic residues provide the protein with the ability to harness the PMF or sense environmental pH in order to promote lipid binding and/or transport. However, more detailed mechanistic studies are needed to tease apart the energetics and dynamics of DedA function.

The identification of this putative lipid-binding site raises the question of what role lipid binding plays in YqjA function. UptA from *B. subtilis* and PA4029 bind phospholipids including PE and PG in addition to their preferred UndP substrate^22,25^, suggesting some flexibility in lipid recognition within the DedA family. While YqjA has not been implicated in UndP transport, deletion of *yqjA* has been reported to alter the phospholipid composition of *E. coli*, decreasing PE while increasing PG and cardiolipin levels^42^. These observations make a role for YqjA in phospholipid transport very plausible. However, our data demonstrate lipid interaction rather than transport allowing for other possible interpretations. Previous genetic and physiological studies have linked YqjA to proton motive force maintenance and adaptation to alkaline conditions that are consistent with H^+^ flux activity. Therefore, another possibility is that the bound lipid is not a transported substrate at all and instead plays a structural or catalytic role linked to H^+^ transport activity. As H^+^ flux has not been directly observed for any DedA so far, the link between YqjA function and PMF could also arise indirectly, with the altered lipid composition of the *ΔyqjA* strain having a knock-on effect on proteins involved in PMF homeostasis. An indirect effect of YqjA disruption on respiratory complexes has been postulated previously^53^, although not via lipid composition alteration. Notably, depletion of PE in *E. coli* membrane substantially impairs NADH-dependent respiratory activity and proton gradient generation^54^, thus providing a potential mechanism linking YqjA-dependent lipid homeostasis to PMF maintenance.

In addition to the well conserved functionally important residues, our models also identified Ser52 as a previously unrecognised component of the putative lipid-binding site. Although only modestly conserved at this position across the DedA family^43^, Ser52 is predicted to interact directly with the phospholipid phosphate group in both POPE and POPG models (Fig. 5a,b). Our data reveal that S52C is unable to rescue the alkaline-sensitive phenotype of *ΔyqjA* (Fig. 5h), suggesting that this residue makes a functionally important contribution to YqjA. However, interestingly, the S52C mutant retains the ability to rescue the temperature-sensitive phenotype of the *ΔyqjAΔyghB* mutant (SI Fig. 6), indicating that this mutation does not simply abolish YqjA function. An intriguing possibility is that YqjA possesses two separable activities, lipid transport and H⁺ flux, and that S52 substitution selectively disrupts the latter. In this scenario, the ability of the Ser52 mutant to rescue the temperature-sensitive phenotype could reflect retention of lipid transport activity, whereas its inability to rescue growth under alkaline conditions could result from impaired H⁺ transport. Alternatively, the S52C mutant may simply retain partial activity that is sufficient to complement one phenotype but not the other. Distinguishing between these possibilities will require direct measurement of lipid and H⁺ transport by YqjA and the Ser52 variant.

Overall, this work provides a structural and functional framework for YqjA, in which conserved re-entrant hairpins form the core of a dynamic dimer interface and contribute to a putative phospholipid-binding site. Determining how lipid interaction, conformational dynamics and H⁺ flux are mechanistically linked will be key to unravelling YqjA function and understanding how it influences bacterial physiology.

## Methods

### Molecular biology

All site-directed mutagenesis was performed using the QuikChange II Site-directed mutagenesis kit (Agilent) and the accuracy of all mutant constructs was sequence verified prior to use.

### Protein expression and purification

To overexpress and purify *E. coli* YqjA, the gene encoding YqjA was expressed in *E. coli* C43 cells from a modified pBAD vector containing the *yqjA* gene in-frame with a C-terminal octa-histidine tag (pBADhisYqjA). C43 (DE3) pBADhisYqjA was grown in LB media supplemented with 100 µg/ml ampicillin and incubated at 37 °C at 190 RPM until an OD_600_ of ∼0.7-1.1 was achieved. At this point, expression was induced by addition of L-arabinose to a final concentration of 0.1% (w/v). The cells were then further incubated for a further 2 hours at 37°C at 190 RPM before being harvest by centrifugation 5800 rcf for 15 min. The cell pellet was resuspended in Lysis Buffer (50 mM Tris, pH 7.5, 200 mM NaCl, 5% (v/v) glycerol) supplemented with protease inhibitors, and cells were lysed by 3 passe through a cell disruptor (Avestin Emulsiflex C3). The lysate was clarified by centrifugation at 20,000 rcf 20 min at 4°C. The supernatant was collected and ultracentrifuged at 200,000 rcf for 90 minutes. The membrane pellet was resuspended in Purification Buffer (20 mM Tris, pH 7.5, 200 mM NaCl, 5% (v/v) glycerol), flash frozen using liquid nitrogen and stored at -80 °C. YqjA-containing membrane fraction was solubilised by incubation with 3.92 mM *n*-Dodecyl-β-D-maltoside (DDM) for 90 min at 4°C. The insoluble fraction was removed by ultracentrifuged at 200,000 rcf for 30 min at 4 °C, and the soluble fraction was incubated Talon resin (Takara) and incubated with agitation for 90 min at 4 °C. The protein/resin mix decanted into a gravity flow column, the flowthrough was collected, the resin was washed with PB supplemented with 10 mM imidazole and 0.2 mM DDM. Bound protein was eluted with PB containing 300 mM imidazole and 0.2 mM DDM and further purified using size exclusion chromatography (SEC) with a Superdex 200 Increase 10/300 GL column (GE Healthcare) and SEC buffer (50 mM Tris, pH 7.5, 400 mM NaCl, 5% (v/v) glycerol, 0.24 mM DDM).

### Analytical ultracentrifugation (AUC)

Sedimentation velocity experiments were performed on YqjA at four different concentrations, 0.59, 1.3, 2.6 and 5.9 mg/ml, in SEC buffer. The samples were ultracentrifuged at 130,000 rcf at 4 °C using a Beckman XL-I analytical ultracentrifuge with an An-50 Ti rotor (Beckman Coulter, USA). Samples of 100 or 50 µL were loaded into two-channel titanium centerpieces of 3- or 1.5-mm optical path-lengths, equipped with sapphire windows (Nanolytics, Germany). The buffer in the reference channel was the same as the proteins but devoid of detergent. The acquisition was done every 8 min, overnight, using absorbance at 280 nm and interference detection.

Data processing and analysis were performed using SEDFIT software v 16.36^55^, REDATE v 1.0.1_56_ and GUSSI v 1.4.2^57^, using standard equations and protocols described previously^37,58^. Sedimentation velocity profiles were globally analysed in terms of *c*(*s*) distributions. The partial specific volume, refractive index increment (*dn/dc*), and extinction coefficient at 280 nm of YqjA (0.753 ml/g, 0.193 ml/g and 1.351 ml/(mg cm), respectively) were estimated, at 4 °C, from the amino-acid sequence, using SEDFIT software v 16.36. The partial specific volume and *dn/dc* of DDM are 0.82 ml/g and 0.143 ml/g, respectively^59^. We considered that DDM does not absorb at 280 nm. The solvent density (ρ = 1.030 g/ml) and the solvent viscosity (η = 1.852 mPa s) were estimated using Sednterp software v 3.0.2 Beta. *c*(*s*) distributions and corrected *s*-values, *s*_20w_, were calculated, considering for the partial specific volume, the mean value between protein and detergent. The amount of bound detergent was calculated from the integration of the *c*(*s*) signals at 280 nm and in interference, using GUSSI^37^. Derivation of the association state from the *s*-value was done considering a globular compact shape (frictional ratio of 1.25), a theoretical molar mass of YqjA monomer of 26.5 kDa, and the appropriate amount of bound detergent.

### Protein structure modelling

To model the YqjA dimer, the the full-length YqjA sequence from *E. coli* (UniProt: P0AA63) was used as input to AlphaFold-multimer v2.2^39,40^, v2.3 (https://github.com/google-deepmind/alphafold/blob/main/docs/technical_note_v2.3.0.md) and AFsample^60^ and AlphaFold 3^61^. The full sequence database was used for all the runs. 2,500 and 500 predictions we generated for AlphaFold v.2.2 and v2.3, respectively. Six runs of AFsample were performed with different options (as shown in SI Table 1), amounting to 800 structural predictions. Four runs, each with 1000 predictions, were performed with AlphaFold 3: one with standard parameters, another excluding template information, another adding one 1-palmitoyl-2-oleoyl-sn-glycero-3-phosphoethanolamine (POPE) molecule, and another with a 1-palmitoyl-2-oleoyl-sn-glycero-3-phospho-(1’-rac-glycerol) (POPG) molecule. The models were ranked using the AlphaFold-Multimer score, defined as a combination of the predicted template modelling (pTM) and predicted interface template modelling (ipTM) scores (0.2pTM + 0.8ipTM). Predicted dimer structures that have a value of 0.2 or 0.4 and above, depending on the sampled scores, were filtered. We chose a minimum score of 0.2 because values below this threshold may also reflect low confidence in the protomer structure prediction.

The contacts between each protomer of the dimer structural predictions were then counted. A contact was computed if any heavy atom of one residue was within 4.2 Å of any heavy atom of another residue. The minimum distance was measured between the topological segments (as defined previously^32^) that formed frequent contacts in the predicted dimers. To identify the main interfaces sampled, they were clustered in the space of this minimum distance using the algorithm of Daura^62^ coded in the METAGUI plugin^63^.

### Site-specific crosslinking

Copper phenanthroline-based crosslinking was adapted from a previous study^32^. Briefly, for whole cell chemical crosslinking, the expression strain C43 pBADhisYqjA (or mutant variant), a single colony of transformed cells was grown overnight at 37 °C in LB supplemented with 100 µg/ml ampicillin. Following overnight incubation, the cultures were diluted to an OD_600_ of 0.2 using fresh LB supplemented with 100 µg/µl ampicillin and grown at 30°C for 90 min. Protein expression was then induced by addition of 0.1% (w/v) L-arabinose, and the cells were incubated for a further 60 minutes at 30 °C with agitation. The cells were harvested by centrifugation at 4000 rcf and then resuspended in Purification buffer to an OD_600_ of 0.8-1.0. The resuspended cells were divided into three aliquot and were incubated for 30 min at 20°C with either 5 mM dithiothreitol (DTT), 0.6 mM copper phenanthroline (CuPhen) or no addition (buffer was added to ensure equal reaction volume in all samples). Crosslinking was quenched by addition of 100 mM of methyl-methanethiosulfonate (MMTS). Samples were visualised using Western blotting using a Tetra-His antibody (Qiagen) and Anti-Mouse IgG peroxidase conjugate secondary antibody (Sigma-Aldrich).

### Phenotypic rescue assays

W3110Δ*yqjAΔyghB* cells were made chemically competent and transformed with pBAD expression vector encoding a variant of *yqjA* or negative controls *gltph* (a non-DedA integral membrane protein) and *siaP* (a soluble sialic acid binding protein). 5 mL of all transformed strains were grown in LB supplemented with 100 µg/mL carbenicillin and incubated overnight at 30 °C. Resulting cultures were normalised to OD_600_ of 1 and serially diluted (100-10 -7x) in in a 96-well plate with LB. 2.5 µl of each dilution was pipetted onto rectangular petri dishes of LB agar supplemented with 0.002% (w/v) L-arabinose. Each of the petri dishes were then incubated at 30 °C and 42 °C for 15 hours.

For growth assays in liquid media, BW25113Δ*yqjA* cells were transformed with a pBAD expression vector encoding a variant of YqjA or a negative control SiaP (sialic acid specific substrate binding protein^64^). Both transformed and untransformed strains were grown overnight at 37°C in LB supplemented with 50 µg/mL ampicillin as appropriate. Overnight cultures were subcultured at 37°C to OD600 of 0.5 in LB supplemented with 0.001% L-Arabinose and 50 µg/mL ampicillin, where appropriate and diluted in the same media to an OD600 of 0.5 for the test culture. The pH of the test culture media was adjusted through the addition of 100mM Bis-Tris-Propane and HCl. Growth of test cultures was monitored at 37oC with 300 rpm shaking in a BMG Labtech SPECTROstar Nano.

### GFP thermal stability assay (GFP-TS)

For the GFP-TS assay, YqjA-containing membranes were prepared as previously described for overexpression and purification, except the expression plasmid encoded *yqjA* (or mutant variants) upstream and in-frame with *gfp* and an octahistidine tag. The resultant membranes were solubilised with 3.92 mM DDM and incubated at 4°C for 90 min. The solubilised sample was diluted 27x into GFP-TS buffer (20 mM Tris, pH 7.5, 50 mM NaCl, 1.96 mM DDM and 3.42 mM n-Octyl-ß-D-Glucopyranoside (ß-OG)) and incubated on ice for 15 min. For a full melt curve, membrane samples were incubated at 4, 20, 30, 40, 50, 60 and 70°C for 10 min then centrifuged at 35,000 rcf for 30 min at 4°C. The supernatant was removed and retained, and the remaining pellet was resuspended in GFP-TS buffer to a volume equal to the supernatant fraction. 100 µl of both supernatant and resuspended pellet were aliquoted into a 96 well plate and the GFP fluorescence was measured using a BMG LabTech CLARIOstar plate reader. Proportional melt curves were generated from the fluorescence data using GraphPad Prism, and the was T_m_ determined by the crossover point of supernatant and pellet fluorescence curves.

*E. coli* Polar lipid extracts (PLE) (Avanti) were resuspended with the use of an ultrasonic bath in 20 mM Tris pH 7.5, aliquoted at 100 mg/ml, flash frozen and stored at -80 °C. To test the stabilisation of YqjA in the presence of lipids, samples were incubated in the absence or presence of 5 mg/ml of PLE at a temperature corresponding to the calculated Tm +5 °C for 10 minutes. The data was processed in the same manner as the full melt curve experiments but with only the supernatant proportional GFP fluorescence values being plotted.

## Data availability statement

All the data are contained in the manuscript.

## Author contributions

**Thomas J. Paige:** Investigation, Formal analysis, Writing – original draft, Writing – review & editing. **Junxi Chen:** Methodology, Formal analysis. **Aline Le Roy:** Investigation, Formal analysis, Writing – review & editing. **Connor D. D. Sampson:** Investigation, Writing – review & editing. **Benjamin F. Cooper:** Investigation, Writing – review & editing. **Matthew Batson:** Investigation, Writing – review & editing. **Hollie L. Scarsbrook:** Investigation. **Lucy R. Forrest:** Supervision, Formal analysis, Methodology, Resources, Writing – review & editing. **Georgia L. Isom:** Supervision, Formal analysis, Resources, Writing – review & editing. **Caroline Mas:** Supervision, Formal analysis, Methodology, Writing – review & editing. **Vanessa Leone:** Methodology, Formal analysis, Funding acquisition, Resources, Writing – review & editing. **Christopher Mulligan:** Conceptualization, Formal analysis, Funding acquisition, Project administration, Resources, Supervision, Visualization, Writing – original draft, Writing – review & editing.

## Acknowledgements

This work was financially supported by a Biotechnology and Biological Sciences Research Council (BBSRC) grant (UKRI2967), Royal Society Research Grant (RG170265), and Royal Society International Exchange Award (IES\R3\193132) awarded to CM, and the SoCoBio BBSRC doctoral training partnership (BB/T008768/1). This work used the platforms of the Grenoble Instruct-ERIC centre (ISBG ; UAR 3518 CNRS-CEA-UGA-EMBL) within the Grenoble Partnership for Structural Biology (PSB), supported by FRISBI (ANR-10-INBS-0005-02) and GRAL, financed within the University Grenoble Alpes graduate school (Ecoles Universitaires de Recherche) CBH-EUR-GS (ANR-17-EURE-0003). This research was completed in part with computational resources and technical support provided by the Research Computing Center at the Medical College of Wisconsin.

## Supplementary information

**SI Table 1:** used AFsample options.

| **SI Table 1: used AFsample options** |  |  |  |  |  |  |
| --- | --- | --- | --- | --- | --- | --- |
| Run number | Number of structures | Maximum recycles | Dropout? | Alphafold multimer model | Dropout structure module? | Template |
| 1 | 100 | 3 | False | Multimer_v1 | False | Yes |
| 2 | 100 | 3 | False | Multimer_v2 | False | Yes |
| 3 | 200 | 3 | True | Multimer_all | True | Yes |
| 4 | 100 | 3 | True | Multimer_v1 | False | Yes |
| 5 | 100 | 21 | True | Multimer_v1 | False | Yes |
| 6 | 200 | 3 | True | Multimer_v2 | False | Yes |

**Supplementary figure 1.**
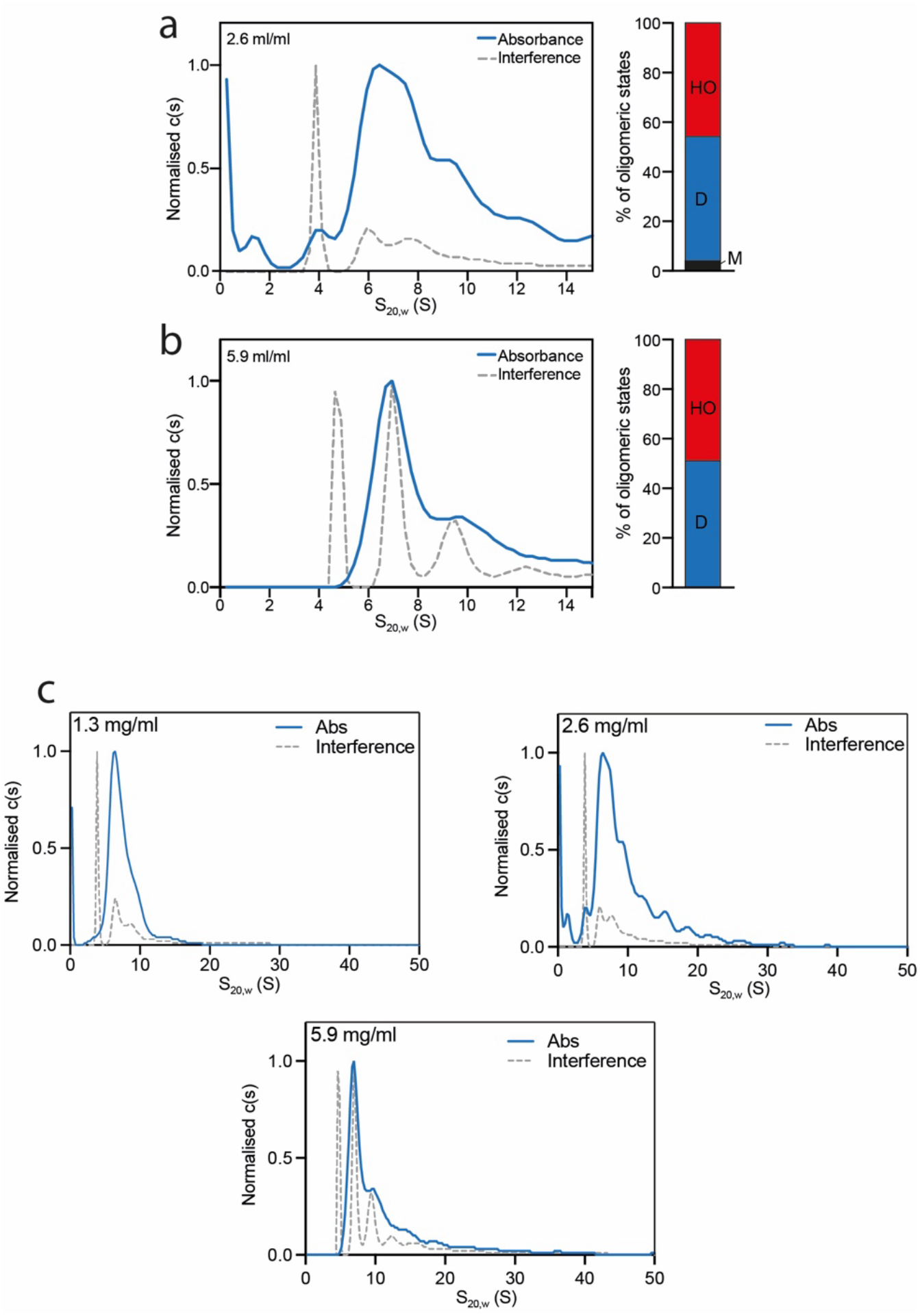
Analytical ultracentrifugation (AUC) analysis of YqjA. Normalised c(s) distributions obtained at 280 nm (continuous blue line) and using interference detection (dashed grey lines), for YqjA at **a)** 2.6 mg/ml and **b)** 5.9 mg/ml. **c)** Normalised c(s) distributions obtained at 280 nm (continuous blue line) and using interference detection (dashed grey lines), for YqjA at 1.3, 2.6 and 5.9 mg/ml with expanded x-axis to observe the high S_20w_ values.

**Supplementary figure 2.**
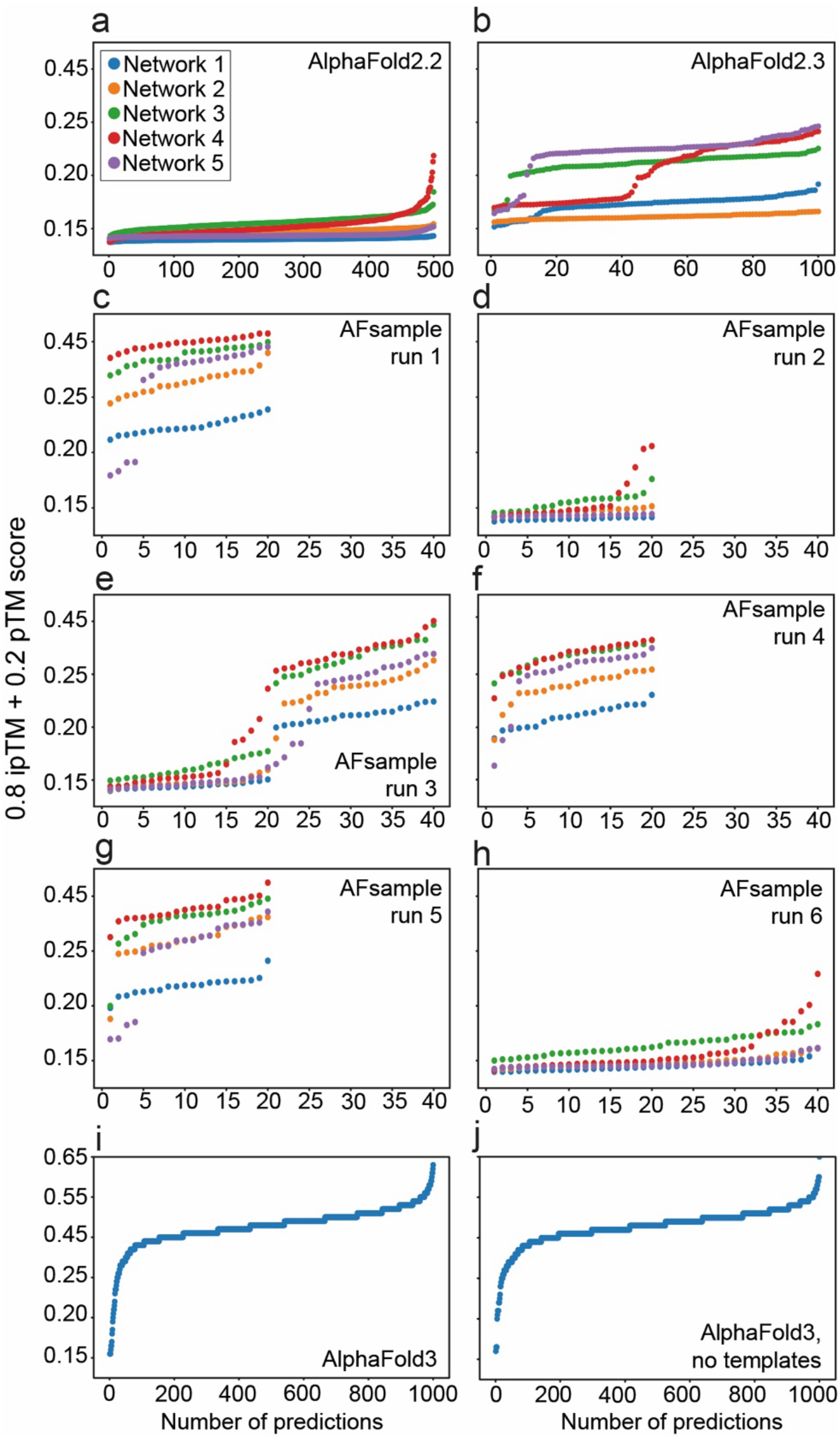
AlphaFold multimer scores. Sorted combined pTM and ipTM scores per predicted structure by each neural network in all the modeling runs.

**Supplementary Figure 3.**
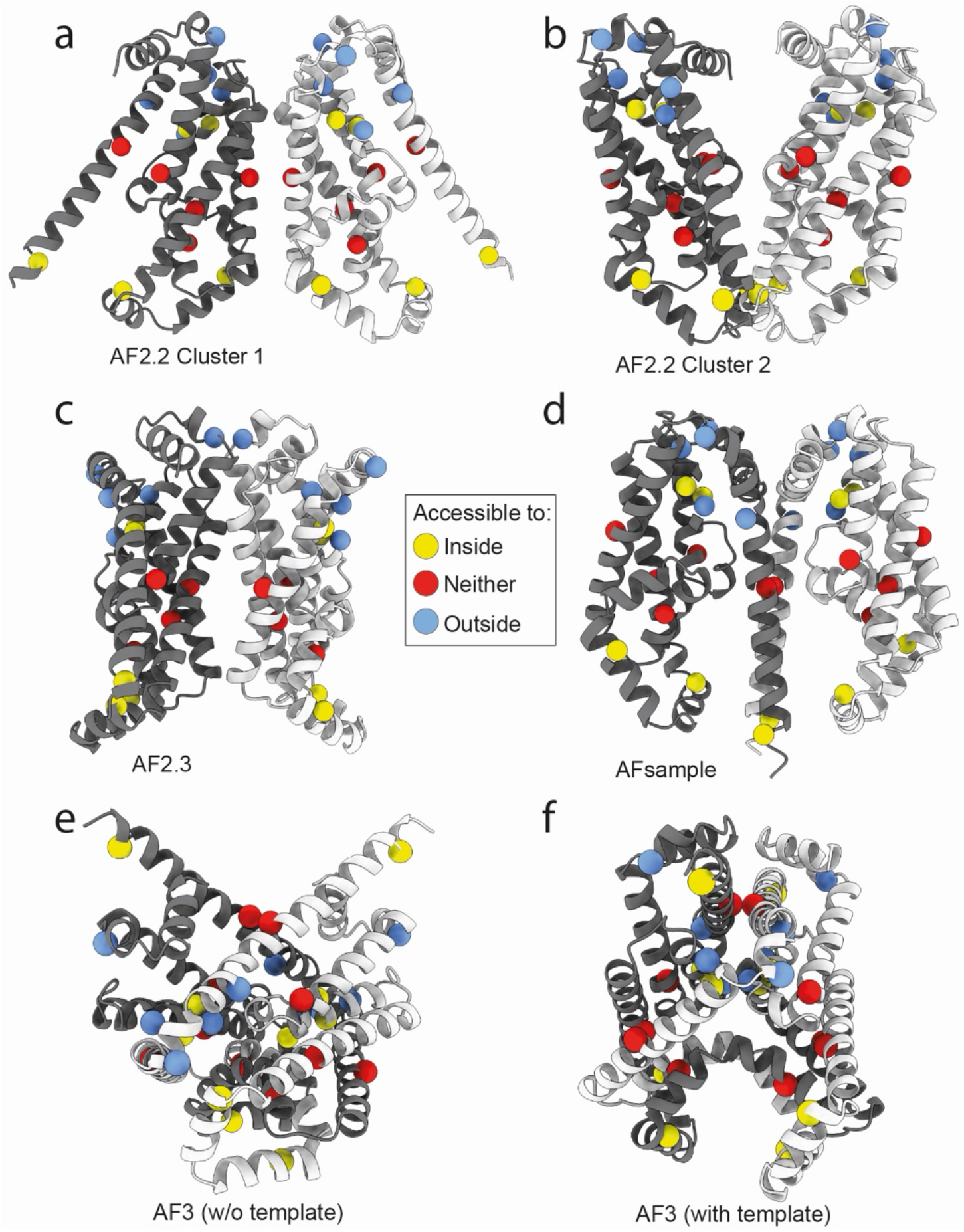
Selected structures per each modeling run and compatibility with previous SCAM data. The final structures were selected clustering in the space of the minimum distances between transmembrane regions, as explained in the Methods. One protomer is dark grey and the other is coloured white. The carbon alpha atoms of the residues assessed with cysteine accessibility^32^ are shown as balls. They are colored depending on the whether the SCAM data indicated these positions were accessible from the outside (blue, reacted with both MTSES and NEM), the inside of the cell (yellow, reacted with only membrane permeable NEM) or accessible to neither (red, reacted with neither cysteine-reactive reagent)^32^.

**Supplementary figure 4.**
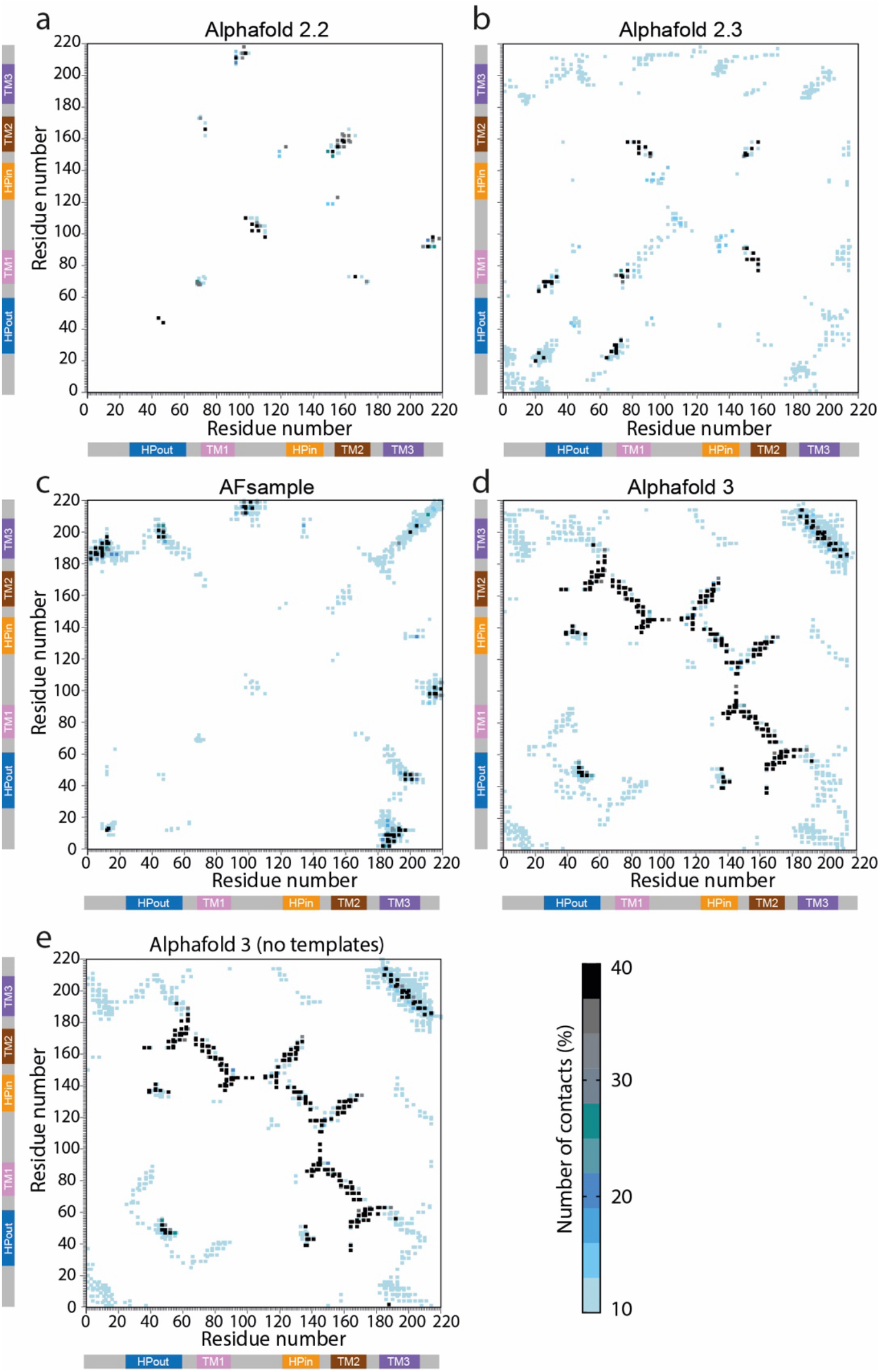
Interprotomeric interactions. Pairwise interaction between the protomers of YqjA predicted dimers after filtering based on the combined ipTM and pTM score (prediction with a score above 0.2) from the different modelling approaches; **a)** AF2.2, **b)** AF2.3, **c)** AFsample, **d)** AF3 (with templates), **e)** AF3 (without templates). Two residues are defined as interacting if any non-hydrogen atom of one of the residues is at 4.2 Å of any non-hydrogen atom of the other residue. Each pairwise contact is colored based on the number of times that it has been observed in the structural predictions.

**Supplementary figure 5.**
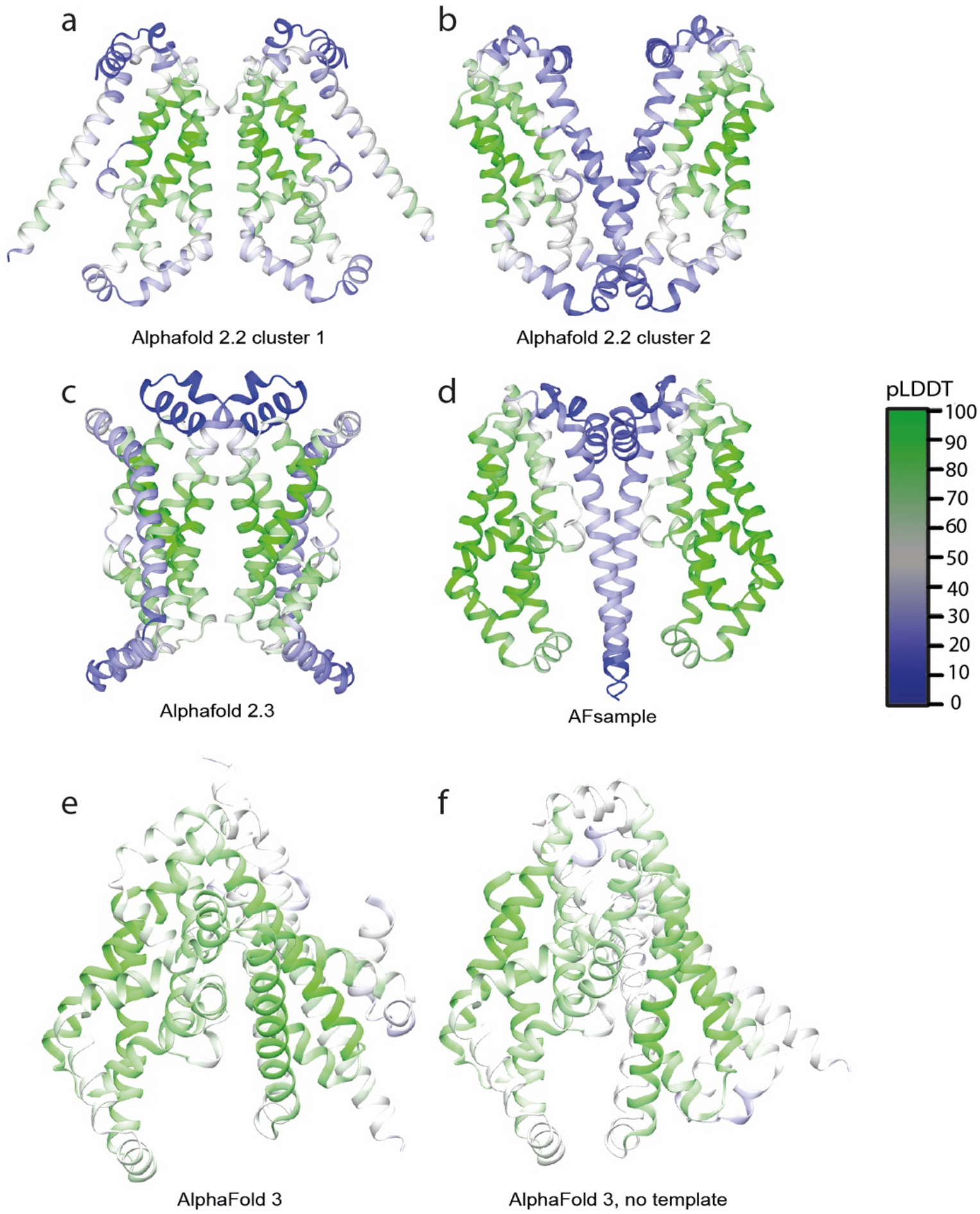
**Per residue estimate of structural confidence**. pLDDT score of the selected structural predictions of each run. **a)** AF2.2, cluster 1, **b)** AF2.2, cluster 2, **c)** AF2.3, **d)** AFsample, **e)** AF3 (with templates), **f)** AF3 (without templates).

**Supplementary figure 6.**
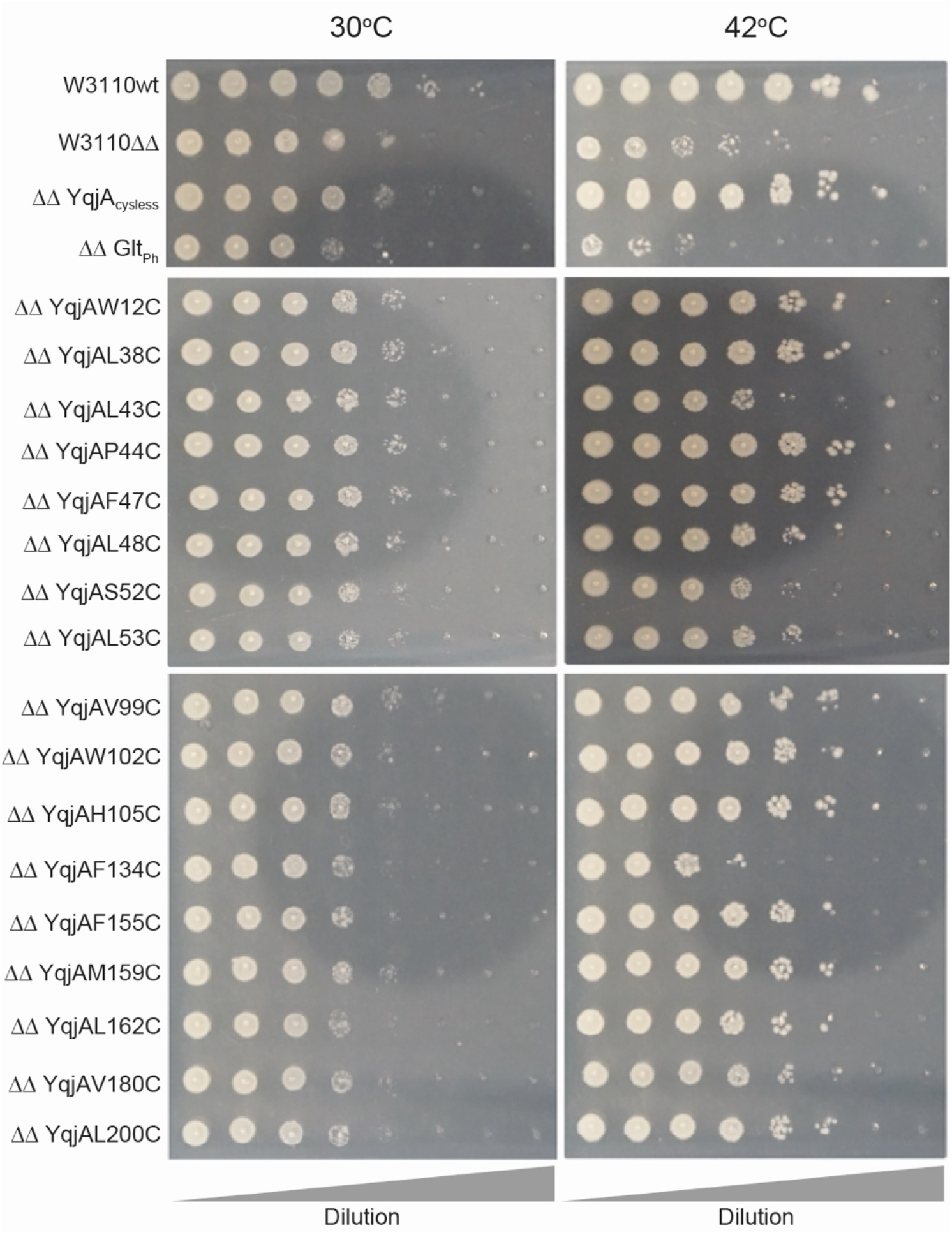
Phenotype rescue assay to assess function of YqjA cysteine mutants. Phenotypic rescue assays wherein wildtype *E. coli* W3110 (W3110wt) or W3110 Δ*yqjA,ΔyghB* (W3110ΔΔ/ΔΔ) cells harbouring plasmids expressing cysteine-free *yqjA* (YqjA_cysless_), *yqjA* cysteine variants or control plasmid (Glt_Ph_). Cells were grown on LB agar with 0.002% arabinose and incubated at either 30 °C or 42 °C for 15 hours. The assays were repeated in triplicate and each result was consistent.

**Supplementary figure 7.**
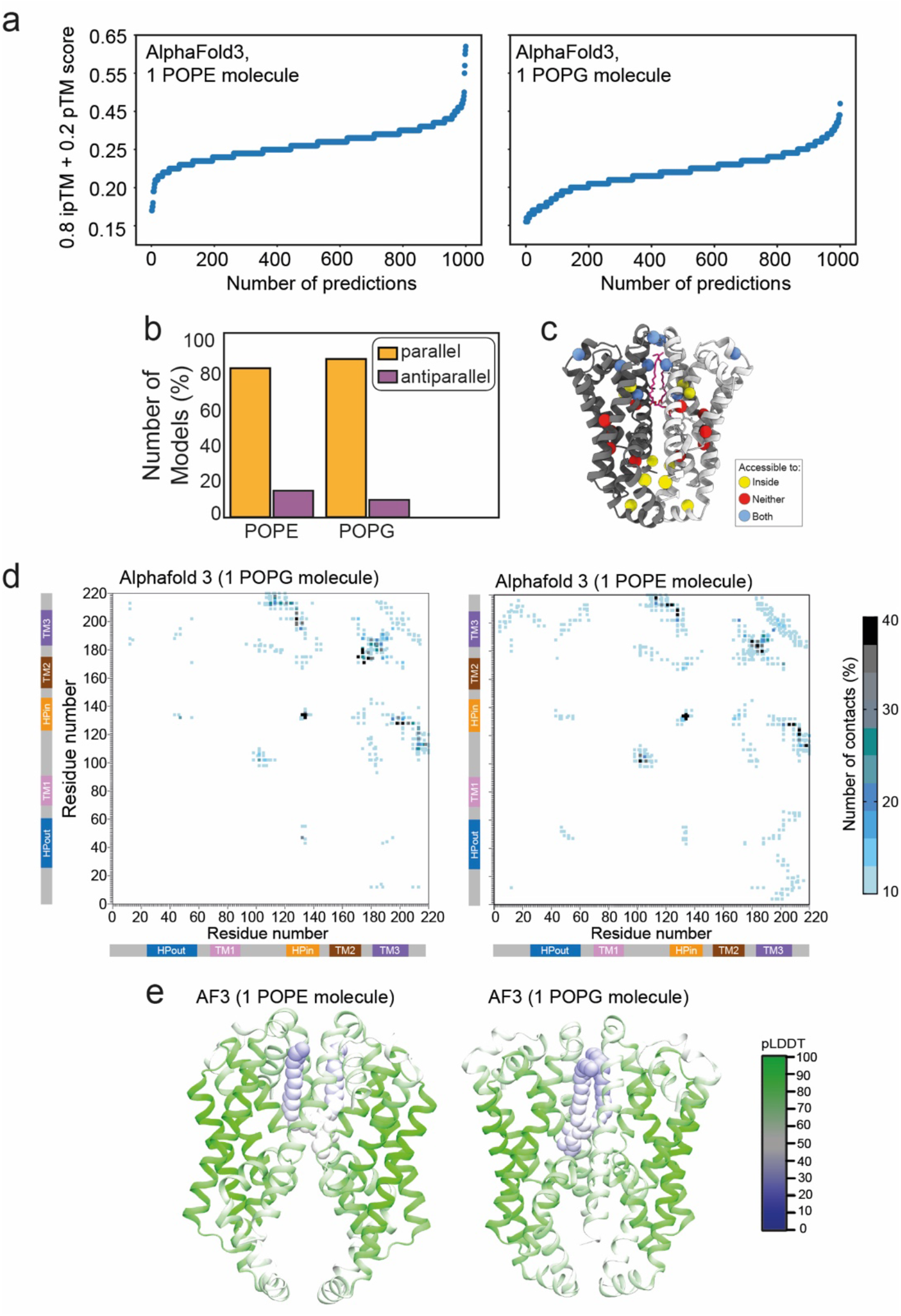
Evaluation of lipid-bound AF3 modelling. **a)** Sorted combined pTM and ipTM scores per predicted structure by AF3 with 1 molecule of POPE (left) or one molecule of POPG (right). **b)** Proportion of parallel and antiparallel complex arrangement. After clustering based on the possible interface (see Methods), clusters were grouped into parallel and antiparallel complexes for the YqjA dimer with a lipid molecule (POPE or POPG). **c)** Cysteine accessibility results from previous SCAM analysis mapped onto AF3, POPE-bound model. Colour coded as in SI Fig. 3. **d)** Contact map showing pairwise interaction between the protomers of YqjA predicted dimers after filtering based on the combined ipTM and pTM score (prediction with a score above 0.2) from AF3 with POPG (left) or POPE (right). **e)** pLDDT score of the selected structural predictions of each run: POPE (left) and POPG (right).

**Supplementary figure 8.**
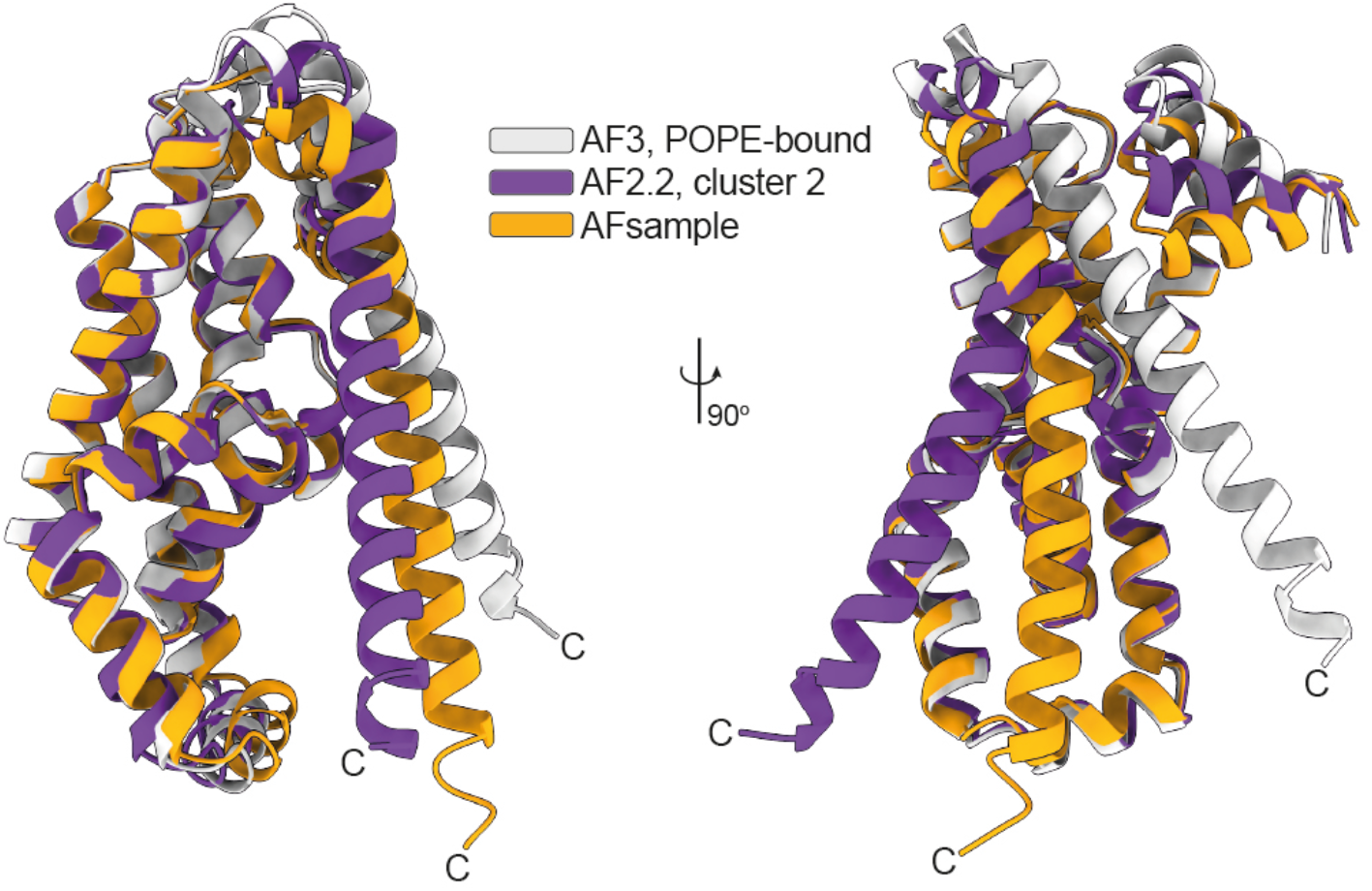
Structural comparison of single protomers from modelling approaches. Superposition of single protomers derived from AF3+POPE (white model), AF2.2, cluster 2 (purple model), AFsample (orange model). C-terminus is labelled in model.

**Supplementary figure 9.**
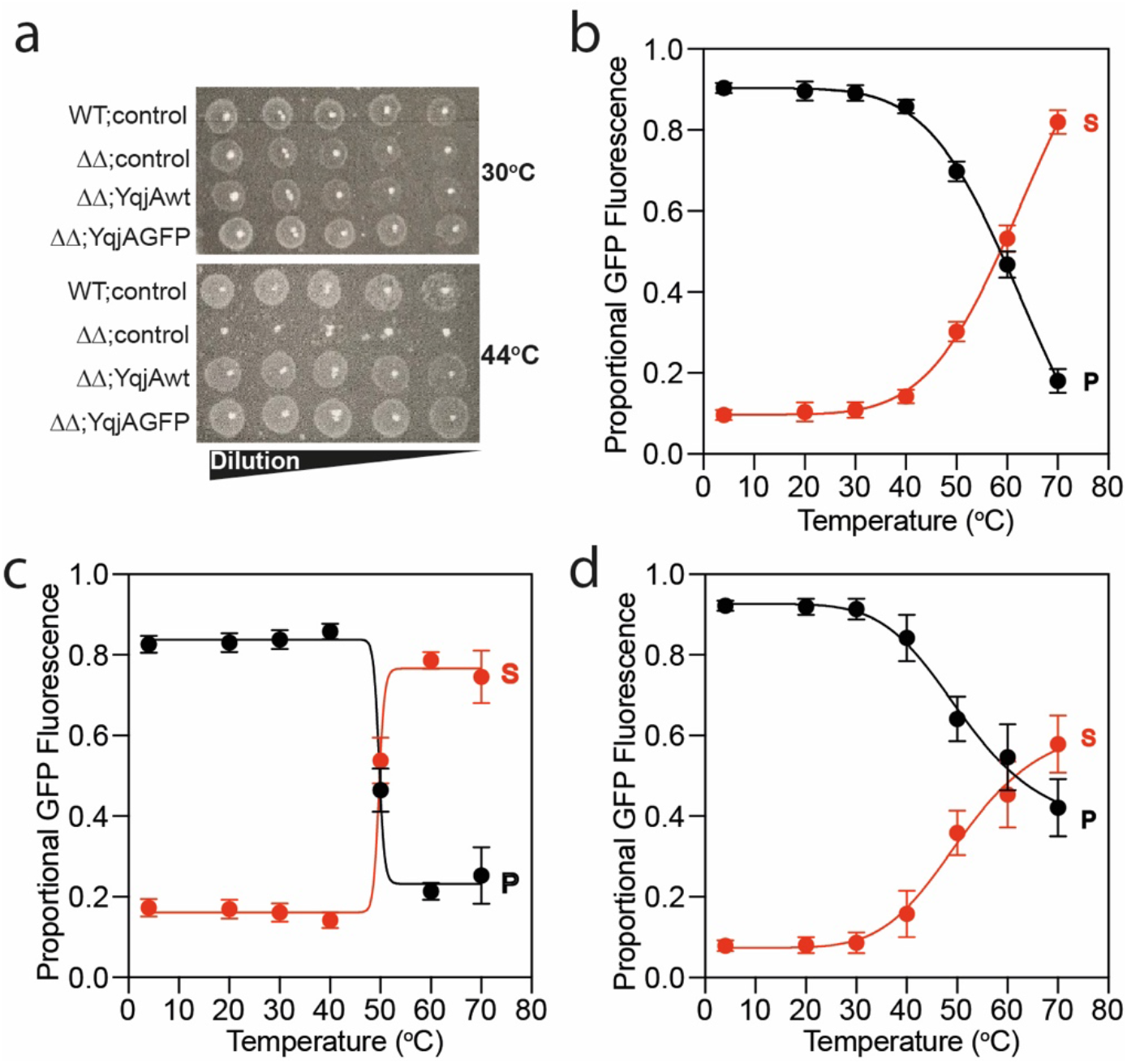
GFP-TS assay with YqjA variants. **a)** Phenotype rescue assay in which wildtype *E. coli* BW25113 (WT) or BW25113Δ*yqjA,ΔyghB* (ΔΔ) cells harbouring plasmids expressing wildtype *yqjA* (YqjAwt), GFP-tagged *yqjA* YqjAGFP) or control plasmid (control, gene encoding aspartate transporter, Glt_Ph_). Cells were grown on LB agar with 0.001% arabinose and incubated at either 30 °C or 42 °C for 20 hours. **b-d)** Proportion of GFP fluorescence in the pellet (black data) and supernatant (red data) as a function of incubation temperature for **b)** E39A/D51A mutant, c) YqjA wildtype in 25 mM NaCl, and d) R130A/R136A in 25 mM NaCl. The mean is plotted (n = 9 for panels b and e, n = 12 for panel c) and error bars represent the standard deviation.

## References

1. Doerrler, W. T., Sikdar, R., Kumar, S. & Boughner, L. A. New functions for the ancient DedA membrane protein family. J. Bacteriol. 195, 3–11 (2013).

2. Liang, F. T. et al. BB0250 of Borrelia burgdorferi is a conserved and essential inner membrane protein required for cell division. J. Bacteriol. 192, 6105–6115 (2010).

3. Arai, M., Liu, L., Fujimoto, T., Setiawan, A. & Kobayashi, M. DedA Protein Relates to Action-Mechanism of Halicyclamine A, a Marine Spongean Macrocyclic Alkaloid, as an Anti-dormant Mycobacterial Substance. Mar. Drugs 9, 984–993 (2011).

4. Panta, P. R. & Doerrler, W. T. A Burkholderia thailandensis DedA Family Membrane Protein Is Required for Proton Motive Force Dependent Lipid A Modification. Front. Microbiol. 11, 618389 (2020).

5. Panta, P. R. et al. A DedA Family Membrane Protein Is Required for Burkholderia thailandensis Colistin Resistance. Front. Microbiol. 10, 2532 (2019).

6. Tiwari, V. et al. A Klebsiella pneumoniae DedA family membrane protein is required for colistin resistance and for virulence in wax moth larvae. Sci. Rep. 11, 24365 (2021).

7. Iqbal, A. et al. Chemical or Genetic Alteration of Proton Motive Force Results in Loss of Virulence of Burkholderia glumae, the Cause of Rice Bacterial Panicle Blight. Appl. Environ. Microbiol. 87, e0091521 (2021).

8. Huang, L., Feng, Y. & Zong, Z. Heterogeneous resistance to colistin in Enterobacter cloacae complex due to a new small transmembrane protein. J. Antimicrob. Chemother. 74, 2551–2558 (2019).

9. Tzeng, Y.-L. et al. Cationic antimicrobial peptide resistance in Neisseria meningitidis. J. Bacteriol. 187, 5387–5396 (2005).

10. Shi, Y., Cromie, M. J., Hsu, F.-F., Turk, J. & Groisman, E. A. PhoP-regulated Salmonella resistance to the antimicrobial peptides magainin 2 and polymyxin B. Mol. Microbiol. 53, 229–241 (2004).

11. Boughner, L. A. & Doerrler, W. T. Multiple deletions reveal the essentiality of the DedA membrane protein family in Escherichia coli. Microbiol. Read. Engl. 158, 1162– 1171 (2012).

12. Thompkins, K., Chattopadhyay, B., Xiao, Y., Henk, M. C. & Doerrler, W. T. Temperature sensitivity and cell division defects in an Escherichia coli strain with mutations in yghB and yqjA, encoding related and conserved inner membrane proteins. J. Bacteriol. 190, 4489–4500 (2008).

13. Kumar, S. & Doerrler, W. T. YqjA is a Putative Transporter Essential for Alkaline pH Tolerance in Escherichia Coli. FASEB J. Off. Publ. Fed. Am. Soc. Exp. Biol. 30, 861.1-861.1 (2016).

14. Kumar, S., Tiwari, V. & Doerrler, W. T. Cpx-dependent expression of YqjA requires cations at elevated pH. FEMS Microbiol. Lett. 364, (2017).

15. Sikdar, R. & Doerrler, W. T. Inefficient Tat-dependent export of periplasmic amidases in an Escherichia coli strain with mutations in two DedA family genes. J. Bacteriol. 192, 807–818 (2010).

16. Kumar, S. & Doerrler, W. T. Members of the conserved DedA family are likely membrane transporters and are required for drug resistance in Escherichia coli. Antimicrob. Agents Chemother. 58, 923–930 (2014).

17. Kumar, S. & Doerrler, W. T. Members of the conserved DedA family are likely membrane transporters and are required for drug resistance in Escherichia coli. Antimicrob. Agents Chemother. 58, 923–930 (2014).

18. Kumar, S., Bradley, C. L., Mukashyaka, P. & Doerrler, W. T. Identification of essential arginine residues of Escherichia coli DedA/Tvp38 family membrane proteins YqjA and YghB. FEMS Microbiol. Lett. 363, fnw133 (2016).

19. Ghanbarpour, A., Valverde, D. P., Melia, T. J. & Reinisch, K. M. A model for a partnership of lipid transfer proteins and scramblases in membrane expansion and organelle biogenesis. Proc. Natl. Acad. Sci. U. S. A. 118, (2021).

20. Huang, D. et al. TMEM41B acts as an ER scramblase required for lipoprotein biogenesis and lipid homeostasis. Cell Metab. 33, 1655–1670.e8 (2021).

21. Roney, I. J. & Rudner, D. Z. The DedA superfamily member PetA is required for the transbilayer distribution of phosphatidylethanolamine in bacterial membranes. Proc. Natl. Acad. Sci. 120, e2301979120 (2023).

22. Oluwole, A. O., et al. Lipopeptide antibiotics disrupt interactions of undecaprenyl phosphate with UptA. Proc. Natl. Acad. Sci. 121, e2408315121 (2024).

23. Roney, I. J. & Rudner, D. Z. Two broadly conserved families of polyprenyl-phosphate transporters. Nature 613, 729–734 (2023).

24. Sit, B. et al. Undecaprenyl phosphate translocases confer conditional microbial fitness. Nature 613, 721–728 (2023).

25. Sposato, D. et al. The *Pseudomonas aeruginosa* DedA protein PA4029 is an undecaprenyl phosphate flippase important for polymyxin resistance. mBio 17, e02408–25 (2026).

26. Yan, A., Guan, Z. & Raetz, C. R. H. An undecaprenyl phosphate-aminoarabinose flippase required for polymyxin resistance in Escherichia coli. J. Biol. Chem. 282, 36077– 36089 (2007).

27. Trent, M. S. et al. Accumulation of a polyisoprene-linked amino sugar in polymyxin-resistant Salmonella typhimurium and Escherichia coli: structural characterization and transfer to lipid A in the periplasm. J. Biol. Chem. 276, 43132–43144 (2001).

28. Trent, M. S., Ribeiro, A. A., Lin, S., Cotter, R. J. & Raetz, C. R. An inner membrane enzyme in Salmonella and Escherichia coli that transfers 4-amino-4-deoxy-L-arabinose to lipid A: induction on polymyxin-resistant mutants and role of a novel lipid-linked donor. J. Biol. Chem. 276, 43122–43131 (2001).

29. Breazeale, S. D., Ribeiro, A. A., McClerren, A. L. & Raetz, C. R. H. A formyltransferase required for polymyxin resistance in Escherichia coli and the modification of lipid A with 4-Amino-4-deoxy-L-arabinose. Identification and function oF UDP-4-deoxy-4-formamido-L-arabinose. J. Biol. Chem. 280, 14154–14167 (2005).

30. Mesdaghi, S., Murphy, D. L., Sánchez Rodríguez, F., Burgos-Mármol, J. J. & Rigden, D. J. In silico prediction of structure and function for a large family of transmembrane proteins that includes human Tmem41b. F1000Research 9, 1395 (2020).

31. Okawa, F. et al. Evolution and insights into the structure and function of the DedA superfamily containing TMEM41B and VMP1. J. Cell Sci. 134, (2021).

32. Scarsbrook, H. L., Urban, R., Streather, B. R., Moores, A. & Mulligan, C. Topological analysis of a bacterial DedA protein associated with alkaline tolerance and antimicrobial resistance. Microbiol. Read. Engl. 167, (2021).

33. Keller, R., Schleppi, N., Weikum, J. & Schneider, D. Mutational analyses of YqjA, a Tvp38/DedA protein of E. coli. FEBS Lett. 589, 842–848 (2015).

34. Rath, A., Glibowicka, M., Nadeau, V. G., Chen, G. & Deber, C. M. Detergent binding explains anomalous SDS-PAGE migration of membrane proteins. Proc. Natl. Acad. Sci. U. S. A. 106, 1760–1765 (2009).

35. Currie, M. J. et al. Structural and biophysical analysis of a Haemophilus influenzae tripartite ATP-independent periplasmic (TRAP) transporter. eLife 12, RP92307 (2024).

36. Newton-Vesty, M. C. et al. Structural studies and functional engineering of NanX: an anhydro-sialic acid transporter from Escherichia coli. FEBS Open Bio 10.1002/2211-5463.70310 (2026) doi:10.1002/2211-5463.70310.

37. Salvay, A. G., Santamaria, M., le Maire, M. & Ebel, C. Analytical ultracentrifugation sedimentation velocity for the characterization of detergent-solubilized membrane proteins Ca++-ATPase and ExbB. J. Biol. Phys. 33, 399–419 (2007).

38. Carvalho, F. A. O., Carvalho, J. W. P., Alves, F. R. & Tabak, M. pH effect upon HbGp oligomeric stability: characterization of the dissociated species by AUC and DLS studies. Int. J. Biol. Macromol. 59, 333–341 (2013).

39. Evans, R., et al. Protein complex prediction with AlphaFold-Multimer. *bioRxiv* 2021.10.04.463034 (2022) doi:10.1101/2021.10.04.463034.

40. Jumper, J. et al. Highly accurate protein structure prediction with AlphaFold. Nature 596, 583–589 (2021).

41. Mulligan, C. & Mindell, J. A. Pinning Down the Mechanism of Transport: Probing the Structure and Function of Transporters Using Cysteine Cross-Linking and Site-Specific Labeling. in Methods in Enzymology vol. 594 165–202 (Elsevier, 2017).

42. Thompkins, K., Chattopadhyay, B., Xiao, Y., Henk, M. C. & Doerrler, W. T. Temperature sensitivity and cell division defects in an Escherichia coli strain with mutations in yghB and yqjA, encoding related and conserved inner membrane proteins. J. Bacteriol. 190, 4489–4500 (2008).

43. Todor, H., Herrera, N. & Gross, C. A. Three Bacterial DedA Subfamilies with Distinct Functions and Phylogenetic Distribution. mBio 14, e00028–23 (2023).

44. Nji, E., Chatzikyriakidou, Y., Landreh, M. & Drew, D. An engineered thermal-shift screen reveals specific lipid preferences of eukaryotic and prokaryotic membrane proteins. Nat. Commun. 9, 1–12 (2018).

45. Chatzikyriakidou, Y., Ahn, D.-H., Nji, E. & Drew, D. The GFP thermal shift assay for screening ligand and lipid interactions to solute carrier transporters. Nat. Protoc. 16, 5357–5376 (2021).

46. Sampson, C. D. D., Fàbregas Bellavista, C., Stewart, M. J. & Mulligan, C. Thermostability-based binding assays reveal complex interplay of cation, substrate and lipid binding in the bacterial DASS transporter, VcINDY. Biochem. J. 478, 3847–3867 (2021).

47. Hattori, M., Hibbs, R. E. & Gouaux, E. A fluorescence-detection size-exclusion chromatography-based thermostability assay for membrane protein precrystallization screening. Structure 20, 1293–1299 (2012).

48. Kumar, S. & Doerrler, W. T. Escherichia coli YqjA, a Member of the Conserved DedA/Tvp38 Membrane Protein Family, Is a Putative Osmosensing Transporter Required for Growth at Alkaline pH. J. Bacteriol. 197, 2292–2300 (2015).

49. Drew, D. & Boudker, O. Shared Molecular Mechanisms of Membrane Transporters. Annu. Rev. Biochem. 85, 543–572 (2016).

50. Lambert, E., Mehdipour, A. R., Schmidt, A., Hummer, G. & Perez, C. Evidence for a trap-and-flip mechanism in a proton-dependent lipid transporter. Nat. Commun. 13, 1022 (2022).

51. Bergman, S., Cater, R. J., Plante, A., Mancia, F. & Khelashvili, G. Substrate binding-induced conformational transitions in the omega-3 fatty acid transporter MFSD2A. Nat. Commun. 14, 3391 (2023).

52. Cater, R. J. et al. Structural basis of omega-3 fatty acid transport across the blood– brain barrier. Nature 595, 315–319 (2021).

53. Doerrler, W. T., Sikdar, R., Kumar, S. & Boughner, L. A. New functions for the ancient DedA membrane protein family. J. Bacteriol. 195, 3–11 (2013).

54. Mileykovskaya, E. I. & Dowhan, W. Alterations in the electron transfer chain in mutant strains of Escherichia coli lacking phosphatidylethanolamine. J. Biol. Chem. 268, 24824–24831 (1993).

55. Schuck, P. Size-Distribution Analysis of Macromolecules by Sedimentation Velocity Ultracentrifugation and Lamm Equation Modeling. Biophys. J. 78, 1606–1619 (2000).

56. Zhao, H. et al. A Multilaboratory Comparison of Calibration Accuracy and the Performance of External References in Analytical Ultracentrifugation. PLOS ONE 10, e0126420 (2015).

57. Brautigam, C. A. Chapter Five - Calculations and Publication-Quality Illustrations for Analytical Ultracentrifugation Data. in Methods in Enzymology (ed. Cole, J. L.) vol. 562 109–133 (Academic Press, 2015).

58. Roy, A. et al. AUC and Small-Angle Scattering for Membrane Proteins. in Methods in Enzymology vol. 562 (2015).

59. Salvay, A. G. & Ebel, C. Analytical Ultracentrifuge for the Characterization of Detergent in Solution. in Analytical Ultracentrifugation VIII (eds Wandrey, C. & Cölfen, H.) 74–82 (Springer Berlin Heidelberg, Berlin, Heidelberg, 2006).

60. Wallner, B. AFsample: improving multimer prediction with AlphaFold using massive sampling. Bioinformatics 39, btad573 (2023).

61. Abramson, J. et al. Accurate structure prediction of biomolecular interactions with AlphaFold 3. Nature 630, 493–500 (2024).

62. Daura, X., et al. Peptide Folding: When Simulation Meets Experiment. Angew. Chem. Int. Ed. 38, 236–240 (1999).

63. Giorgino, T., Laio, A. & Rodriguez, A. METAGUI 3: A graphical user interface for choosing the collective variables in molecular dynamics simulations. Comput. Phys. Commun. 217, 204–209 (2017).

64. Mulligan, C. et al. The substrate-binding protein imposes directionality on an electrochemical sodium gradient-driven TRAP transporter. Proc. Natl. Acad. Sci. U. S. A. 106, 1778–1783 (2009).

## References

1 Scarsbrook, H. L., Urban, R., Streather, B. R., Moores, A. & Mulligan, C. Topological analysis of a bacterial DedA protein associated with alkaline tolerance and antimicrobial resistance. Microbiol. Read. Engl. 167, (2021).

